# Does testosterone impair men’s cognitive empathy? Evidence from two large-scale randomized controlled trials

**DOI:** 10.1101/516344

**Authors:** Amos Nadler, Colin F. Camerer, David T. Zava, Triana L. Ortiz, Neil V. Watson, Justin M. Carré, Gideon Nave

## Abstract

The capacity to infer others’ mental states (known as “mind reading” and “cognitive empathy”) is essential for social interactions across species, and its impairment characterizes psychopathological conditions such as autism spectrum disorder and schizophrenia. Previous studies reported that testosterone administration impaired cognitive empathy in healthy humans, and that a putative biomarker of prenatal testosterone exposure (finger digit ratios) moderated the effect. However, empirical support for the relationship has relied on small-sample studies with mixed evidence. We investigate the reliability and generalizability of the relationship in two large-scale double-blind placebo-controlled experiments in young men (*N*=243 and *N*=400), using two different testosterone administration protocols. We find no evidence that cognitive empathy is impaired by testosterone administration or associated with digit ratios. With an unprecedented combined sample size, these results counter current theories and previous high-profile reports, and demonstrate that previous investigations of this topic have been statistically underpowered.

---

Decades of research on neuroendocrinological influences on animal behaviour has provided a reliable basis for exploring it in humans [1] and motivate a growing scientific focus on the biological basis of social aptitudes and the causes of their deficits [2]. One important element of social cognition is *“cognitive empathy”*, which constitutes the capacity to infer from observation the emotions, beliefs, and goals of others.^1^ This capacity exists across taxa [3] where its impairment in humans characterizes a broad range of psychopathological conditions and is part of the clinical diagnostic criteria for autism spectrum disorders (ASDs) [4].^2^

## Testosterone-based biological theory of social cognition

A popular biopsychological model known as the Extreme Male Brain (EMB) hypothesis [5] proposes that two distinct cognitive styles, *“systemizing”* and *“empathizing”*, typify males and females respectively. The stereotypically male systemizing domain has no social dimension, and in its extreme form social cognition is extinguished. Guided by observations that ASDs emerge early in life and are substantially more prevalent among males^3^, and that males typically score lower than females in tests of cognitive empathy [6], the EMB hypothesis proposes that elevated prenatal exposure to the sex steroid testosterone causes impairments in cognitive empathy, through its masculinizing effect on the developing brain [7].

The EMB hypothesis found evidential support in a study that reported a correlation between amniotic testosterone levels and ASD traits ([8], though see [9]), and has remained popular yet controversial to date. Much of its research has relied on the assumption that the ratio between the hand’s second (index) to fourth (ring) digit (known as 2D:4D) is a developmental proxy for prenatal testosterone exposure [10], which motivated examinations of correlations between 2D:4D and cognitive empathy and ASDs occurrences. While some studies provided supporting evidence (e.g., [11]), several others failed to detect a relationship between digit ratio and cognitive empathy [12,13]. Moreover, because it is not feasible to experimentally manipulate prenatal testosterone exposure in humans (due to ethical considerations), findings along this line of research have been correlational, which cannot establish causal relations [8].

## Testing testosterone’s causal effect on cognitive empathy

A handful of experiments attempted to address the above limitation by testing the effects of testosterone administration on cognitive empathy in neurotypical adults, and investigating the dependency of these effects on the 2D:4D biomarker [14–17]. This line of research critically relies on an assumption originating in animal research, that *in utero* androgen exposure moderates the activational effect of testosterone [18]. The seminal publication along this line of research reported a placebo-controlled within-subject experiment of 16 healthy females, in which exogenously administered testosterone strongly impaired cognitive empathy measured using the “Reading the Mind in the Eyes Test” (RMET), a 36-item battery testing the ability to infer others’ emotional states and intentions from pictures of their eye regions [6] (see Fig. S2 in Supplementary Material for example item). In addition to reporting a main effect of exogenous testosterone reducing cognitive empathy, more than 50% of the individual differences in the effect on the RMET were explained by the participants’ variation in the right-hand 2D:4D, implying involvement of prenatal testosterone exposure in the causal effect [17].

A similar experiment with roughly twice the sample size (*N*=33, all female sample) found a much smaller^4^ main effect (*P*=0.048, one-tailed), and no moderation by 2D:4D [16]. A third experiment of 16 females found neither a main effect nor a moderation by 2D:4D [15]. Last, one experiment investigated the effect of testosterone administration on the RMET in 30 healthy males and found neither a main effect nor a moderation by the right-hand 2D:4D; however, subsample analysis revealed that testosterone administration reduced cognitive empathy in participants with relatively low (i.e., more masculine) left-hand 2D:4D, but no relationship for high 2D:4D or either for the right hand [14].

## Do the data support the hypothesis?

Despite these earlier findings the current literature on the effects of testosterone on cognitive empathy is subjected to important limitations and results reveal weaknesses under scrutiny. First, albeit there are a few parallel findings in terms of negative direction of effect of testosterone on RMET performance, there is a lack of replicability across experiments, where only one of the four studies observed a statistically significant (P<0.05) main effect of testosterone on the RMET [17]. Moreover, this publication’s report of a strong moderating role of the right-hand 2D:4D was not replicated in any of the other studies (the only other report of an interaction between testosterone administration and the 2D:4D was observed for the left hand [14]).^5^

A second concern is statistical power. Although the RMET is a noisy psychological instrument,^6^ and 2D:4D is, at best, a noisy proxy of prenatal testosterone exposure [19], all samples ranged between 16 and 33 participants, which might have been too meager to credibly estimate a true effect size. It is therefore impossible to know whether the inconsistencies in the literature are due to an absence of a true association, or the result of false negative findings due to low statistical power. Thus, these inconsistent results necessitate clarification through additional studies.

To this end, we conducted a powerful direct test of the activational and developmental effects of testosterone on cognitive empathy by measuring the causal effect of exogenous testosterone and the moderating role of putative prenatal androgenic biomarkers in two studies of healthy young men. Our studies constitute the two largest behavioural testosterone administration experiments conducted to date, with samples that were 15 and 25 times greater than the first study that reported a statistically significant effect of testosterone on the RMET in females [17] and 7 and 12 times greater than the largest experiment in males [14]. In both studies we used a computer-based version of the RMET to test the hypothesis that testosterone administration and its purported developmental biomarkers affect cognitive empathy.

## Methods

### Experiment 1

#### Participants and experimental procedure

Two hundred forty-three males aged 18 to 55 (mean age=23.63, SD=7.22) participated in the study and were mostly private Southern California consortium students from diverse ethnic backgrounds (see Participants section and Table S1a in Supplementary Material). The institutional review boards of Caltech and Claremont Graduate University approved the study, all participants gave informed consent, no adverse events occurred during any experimental session, and no participant or researcher was harmed. All data and materials are available on the Open Science Framework (https://osf.io/hztfe/).

Participants registered by their preferred session dates and were added to cohorts of 13 to 16. They arrived at the lab at 9 a.m., signed informed consent forms, and had both of their hands scanned before being randomly assigned to private cubicles where they completed demographic and mood questionnaires (see Supplementary Material for all independent variables) and provided an initial saliva sample by passive drool. Next, participants proceeded to gel application (further details below), after which they were given printed material containing precautions and instructions prior to dismissal (experimental timeline shown in Fig. 1). All participants returned to the lab at 2 p.m., provided a second saliva sample, and began a battery of tasks that lasted approximately two hours. We did not randomize the order of the behavioral tasks, in similar fashion to previous studies [20], to standardize hormonal measurements (which have diurnal cycles) among participants. Following the experiment, participants completed an exit survey, where they indicated their beliefs about the treatment they had received using a five-point scale.

**Figure 1.**
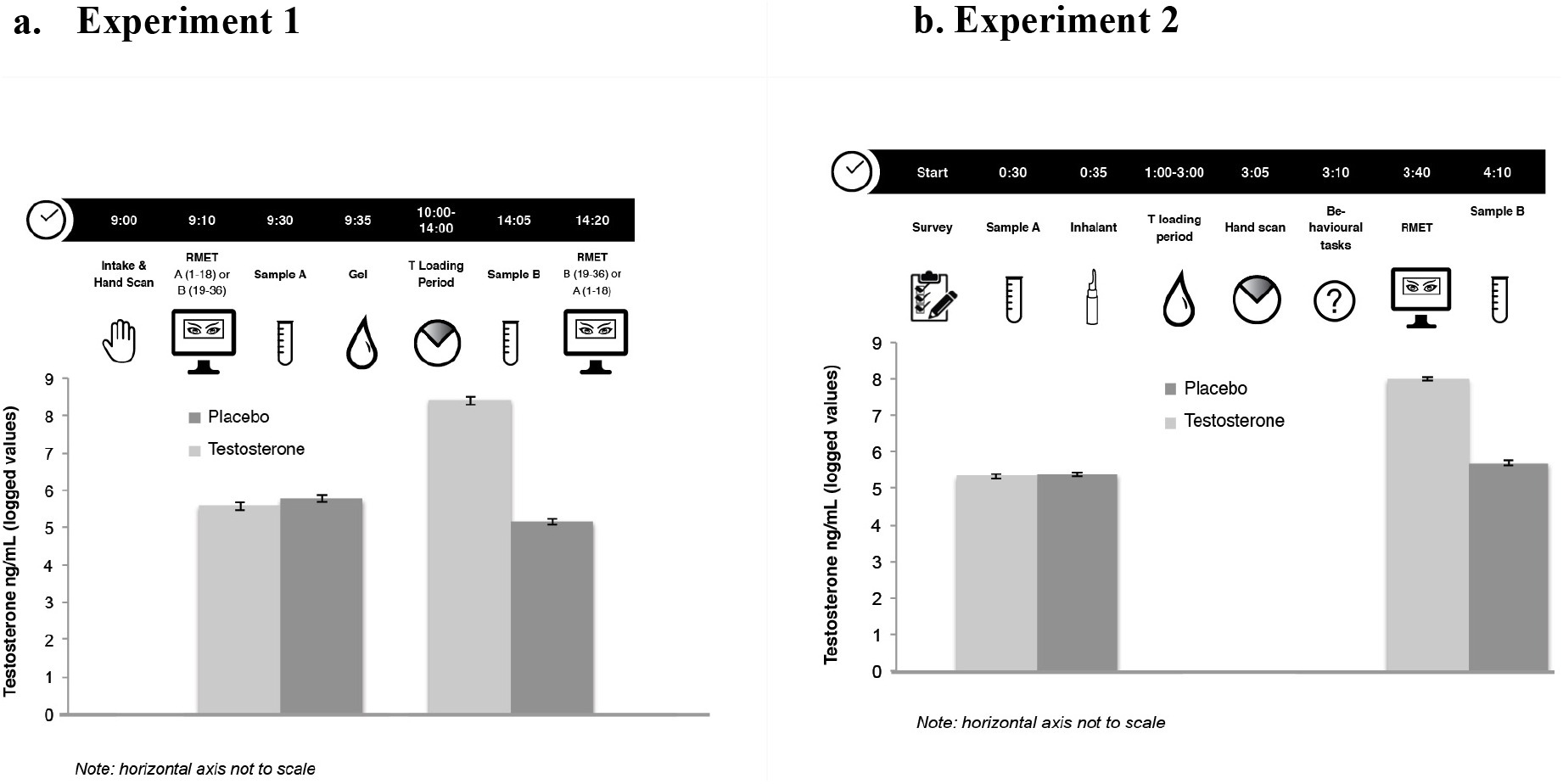
Experimental Timelines. Time-coded experimental timeline shows that, following intake (morning), participants completed half of the RMET (portion A or B), provided a saliva sample, and received gel prior to being dismissed. Upon their return to the lab (afternoon), another saliva sample was collected prior to taking the second portion of the RMET (B or A) (Standard errors shown). Timeline generalizes experimental sequence for all three start times for Experiment 2. Following arrival and completion of consent and self-report questionnaires, participants provided a saliva sample approximately 30 minutes prior to drug administration. The second sample was collected at the end of a two-hour protocol approximately one-and-a-half hours after drug administration and fifteen minutes after the RMET.

#### Treatment administration

Following initial intake, participants were escorted in groups of two to six to a semiprivate room. There they were provided *en masse* small plastic cups containing either 10 g of topical testosterone that is a widely prescribed transdermal testosterone gel with clearly mapped pharmacokinetics [21] (100 mg, Vogelxo™, *N*=123) or volume equivalent of inert placebo of similar viscosity and texture placebo (80% alcogel, 20% Versagel^®^, *N*=118) under a double-blind protocol (see Fig. S1a in Supplementary Material for randomization protocol). Participants were instructed to apply the entirety of the gel container following manufacturer instructions.

#### Saliva samples

Each participant provided four passive drool saliva samples (upon arrival prior to treatment administration, shortly after returning for afternoon session, another closely following the RMET, and a final sample prior to exit survey) for subsequent assay (see Supplemental Material for precise timing). To allow robust manipulation checks and obtain statistical control for hormonal markers of participants’ biological states, we used liquid chromatography tandem mass spectrometry (LC-MS/MS, detection levels and precision are available in Supplementary Material Table S2) to measure the following salivary steroids: estrone, estradiol, estriol, testosterone, androstenedione, DHEA, 5-alpha DHT, progesterone, 17OH-progesterone, 11-deoxycortisol, cortisol, cortisone, and corticosterone (see Supplemental Material Table S7 for measurements).

#### Digit Ratio Measurements

Participants’ hand scans acquired at intake were measured by two independent raters using and digital calipers to quantify 2D:4D; inter-rater correlation was 0.96 and their scores were averaged.

#### Behavioural task

We administered the adult version of the RMET developed by Baron-Cohen and colleagues [6] which shows the eye region of an actor’s face, and a list of four words that describe emotional states and cognitive processes among which participants select the one that best described the person in the image (see Fig. S2 in the Supplemental Material for example item). The task was divided into two segments, baseline (morning) and post-treatment (afternoon), in a repeated measures design (see Fig. 1 below): each participant completed a half of the RMET in the morning (either part “A”, items 1-18, or part “B”, items 19-36) prior to receiving treatment, and the other half following treatment when, according to published pharmacokinetics, androgen levels are elevated and stable following exogenous application.

#### Psychological Questionnaires

Because there are various feasible channels through which testosterone could affect RMET performance (and affect being one of them), we measured mood using the PANAS-X scale [22], both pre- and post-treatment (see Table S1a in Supplementary Material for aggregated responses).

### Experiment 2

#### Participants

Experiment 2 included both students and participants from the general public for a total sample of 400 participants (mean age=22.80, SD=4.68). The all-male sample was composed predominantly of Caucasians and overall ethnic heterogeneity was representative of the region (see Participants section and Table S1b in Supplementary Material). All accepted participants completed the task and were included in the analysis (for pre-screening criteria see Supplementary Material). The Nipissing University Research Ethics Board approved this study, all participants gave informed consent, no adverse events occurred during any experimental session, and no participant or researcher was harmed.

#### Experimental procedure

Participants arrived at one of three testing session times (10:00 a.m., 12:30 p.m., or 2:30 p.m.) in cohorts of six and were brought individually into private testing rooms to read and sign an informed consent form, receive a participant number, and complete questionnaires (see Supplementary Material for all independent variables). Afterwards, participants provided a 1-2 ml saliva sample before treatment administration, after which they had their photos taken and hands scanned. Approximately two hours after arrival to the lab and one-and-a-half hours after drug administration participants completed the RMET then provided their final saliva sample. Upon session completion, participants received compensation and completed an exit survey asking which treatment they believed they had received (see Figure S1b in Supplementary Material).

#### Treatment administration

Following initial saliva sample collection, a researcher provided two syringes pre-filled by a pharmacist following a double-blind protocol each containing 5.5 mg of either placebo or testosterone gel (for a total of 11 mg). This is a newly approved nasal gel used for the treatment of hypogonadism. Pharmacokinetic data indicate that serum testosterone concentrations rise sharply within 15 minutes of testosterone gel application and remain elevated (relative to placebo) up to 180 minutes post-application [23]. The gel was either Natesto^®^ or the volume equivalent of an inert placebo of similar viscosity and texture. Random assignment was determined such that half the participants in every group received testosterone and half received placebo such that total participants base was bisected with *N*=200 for both testosterone and placebo groups (see Fig. S1b in Supplementary Material for randomization protocol).

#### Saliva samples

Each participant provided two saliva samples, with the first sample collection time 30 minutes following arrival (and prior to gel application), and the second 120 minutes after arrival. Participants provided passive drool into a 5 ml polystyrene tube while situated in their individual testing rooms as instructed by a research assistant and samples were analyzed for pre- and post-treatment testosterone and baseline cortisol using commercially available enzyme immunoassay kits (DRG International) (see Table S2b in Supplementary Material for hormone measures).

#### Digit Ratio Measurements

Participants’ 2D:4D were measured by two independent raters using hand scans and digital calipers with an inter-rater correlation of 0.86 and their scores were averaged.

#### Behavioural task

Participants evaluated all 36 items of the RMET [6] as a single task.

### Comparison of experimental features to van Honk et al. (2011)

We note several differences between our study and the primary positive report of testosterone administration on the RMET in 16 females [17].

#### Participant Sex

The EMB theory does not make any sex-specific predictions regarding the developmental and activational effects of testosterone on cognitive empathy. However, van Honk et al. [17] conducted their study in a female-only sample because the pharmacokinetics of a single dose of testosterone had only been studied in females at the time [17]: *“We exclusively recruited women because the parameters … for inducing neurophysiological effects … are known in women but not in men.”* (p. 3450). The recent pharmacokinetic mappings for short-term single dose testosterone administrations [14,21] and availability of two unique administration modalities of FDA-approved exogenous testosterone provided us with a reliable foundation for testing the EMB hypothesis in men.

#### Drug dosage and delivery

van Honk et al. [17] used a sublingual testosterone administration procedure, which causes a sharp increase in serum testosterone of 10-fold or more within 15 minutes, with a rapid decline to normal levels within 90 minutes in women [24]. It is important to note that the pharmacokinetic data for sublingual administration (published in Fig. 1 of Tuiten et al. [24]) show that at the time the task was performed—4 hours after sublingual administration—participants’ testosterone levels were the same across the testosterone and placebo groups. Moreover, the Tuiten et al. study that served as a justification for using a 4-hour delay had only 8 participants, and reported a statistically weak treatment effect (p=0.04, uncorrected for multiple comparisons).

In Experiment 1 we chose to administer testosterone using an FDA-approved transdermal gel for three reasons. First, transdermal gel had been extensively studied in the medical literature both prior and following its approval [25,26]. Second, one of our laboratories found reliable behavioural effects with robust manipulation check in serum using a single dose [27], and third, the pharmacokinetics of a single-dose of this testosterone administration method were mapped prior to the inception of our experiments by a study showing that plasma testosterone levels peaked 3 hours after single-dose exogenous transdermal administration, and stabilized at high levels between 4 and 7 hours following administration [21].^7^ Therefore, we had all participants return to the lab 4.5 hours after receiving gel, when androgen levels were elevated and stable. We used a 100 mg transdermal dose, which quickly elevates then holds testosterone levels high and stable for approximately 24 hours [25] and was shown to generate effects on cognition, decision making, and other behaviours [27,29–31].

In Experiment 2 we used nasal delivery, following a recent study indicating that serum testosterone concentrations rise sharply within 15 minutes after Natesto^®^ gel application and remain elevated for approximately three hours post application among hypogonadal males [23] (see Fig. S3 in Supplementary Material). This method conforms to our experimental paradigm’s pharmacokinetic structure as serum testosterone approached its zenith in treated participants as they completed the RMET (see Supplementary Material Tables S2a and b for testosteron levels).

The doses in both experiments are commonly prescribed daily to men with low circulating testosterone levels and serve as two distinct physical transport channels (transdermal and intranasal, respectively) to reduce the probability that behavioural effects are transport channel specific. Various studies show significant heterogeneity in change in testosterone levels depending on delivery method, location of application in the body, and biofluid measured [14,21,24,27,29,32]. However, all the exogenous delivery methods in this particular literature cause a common hormonal trajectory characterized by a rapid initial rise, a peak above typical circulating levels, and eventual return to baseline.

#### Experimental designs

van Honk et al. [17] used the same questions in pre- and post-treatment testing. As testosterone treatment might affect participants’ capacity to recall answers [33], this design choice might have introduced memory confounds. In Experiment 1 we divided the RMET into two portions, and administered each portion as either pre- or post-treatment measure in, allowing us to capture baseline abilities while ruling out such confounds. In Experiment 2 we conducted a between-subjects experiment that removes all effects of practice and recall from the data.

### Results

#### Manipulation check

In both experiments, pre-treatment (i.e., baseline) saliva testosterone levels were similar across the two treatment groups, and post-treatment saliva testosterone levels were greater in the testosterone group compared with the placebo group. In experiment 1, the mean logged pre-treatment testosterone level was 5.58 pg/ml (*SD*=0.08) in the testosterone group and 5.77 pg/ml (*SD*=0.09) in the placebo group (two-sided t-test: *P*=0.13, *t*(239)=1.50). Mean post-treatment logged testosterone levels were 8.38 pg/ml (*SD*=0.15) in the testosterone group and 5.11 pg/ml (*SD*=0.08) in the placebo group,^8^ (*t*(239)=18.43, *P*<0.0001). Likewise in experiment 2, we find similar mean baseline saliva testosterone concentrations between the groups, with l5.3 pg/ml (*SD*=0.88) in the testosterone group and 5.4 pg/ml (*SD*=0.85) in the placebo group (two-sided t-test: *P*=0.49, *t*(394)=0.69). The mean logged post-treatment saliva testosterone levels were (8.00 pg/ml, *SD*=1.27) in the testosterone group and 5.33 pg/ml (*SD*=0.73) in the placebo group (two-sided t-test: *P*<0.001, *t*(396)=22.14) (see Fig. 1 and Tables S2a and S2b in Supplementary Material).

Consistent with previous reports, we found no treatment effects on mood and treatment expectancy (e.g., [34]) (see Table S2a and S2b in Supplement), or levels of other hormones unaffected by exogenous testosterone, as measured by LC-MS/MS in Experiment 1 or enzyme immunoassay in Experiment 2 (see Tables S1a and S1b in Supplement).

### Influence of Testosterone on RMET scores

Overall RMET scores in our samples were comparable with previous studies of similar populations (see Fig. 2A below). Figure 2B shows baseline and post-treatment RMET scores in Experiment 1, separated by treatment group and order. As expected, baseline (morning) RMET performance were reliably correlated with afternoon scores (*r*(241)=0.40, *P*<0.001). In addition, participants’ scores were, on average, slightly higher in the B portion of the test (A portion average=13.54, *SD*=2.43; B portion average=13.95, *SD*=2.19, *t*(241)=2.53, *P*=0.01). Figure 2C shows Experiment 2 scores by treatment groups.

**Figure 2.**
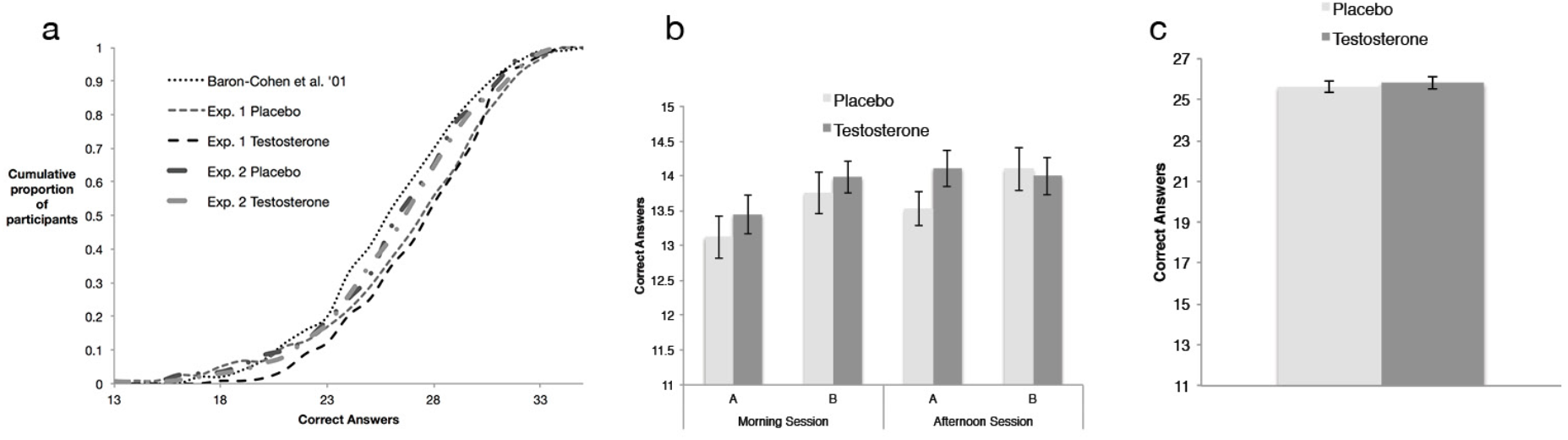
RMET distributions and scores across treatment groups. (*A*) Cumulative distributions of scores from our experiments juxtaposed with the original paper [6]; scores for Experiment 1 portions A and B (both of which are unaffected by treatment) are combined into a total score: RMET scores in Experiment 1 was 27.5 (*SD*=3.9), and 25.6 (*SD*=3.9) in Experiment 2, which are similar to male students in [6] showing average score of 27.3 (*SD*=3.7). (*B*) Experiment 1 pre- and post-treatment RMET scores. No pre- or post-treatment differences were found between the two groups, regardless of the order in which portions of the tests were taken (standard errors shown). (*C*) Experiment 2 RMET scores. No differences were found between treatment groups.

To test for the main effect of testosterone administration on cognitive empathy for Experiment 1, we estimated linear regression models with the post-treatment (afternoon) RMET score as the dependent variable, a binary treatment indicator (testosterone =1, placebo=0) as the independent variable of primary interest, controlling for baseline performance, the order of the two portions of the RMET (A and B) and additional control variables^9^ (the results remain unchanged when these control variables are excluded from the models; see Tables S3a and S4a in Supplementary Material). Analogously, Experiment 2 data were analyzed using linear models with total RMET score as the dependent variable, a binary treatment indicator as the chief independent variable of interest, and control variables^10^ (results remain unchanged with their exclusion from the models; see Supplementary Material Tables S3b and S4b).

We found no reliable effect of testosterone administration on the RMET in Experiment 1 (beta=0.11, 95% CI=[−0.45, 0.68]; *t*(237)=0.37, *P*=0.71; Cohen’s *d*=0.04, 95% CI=[−0.19, 0.28]). Thus, the effect’s point estimate was positive and the 95% CI excluded the *d*=−0.49 reported in (van Honk et al. 2011) or any *negative* effects that are greater in magnitude than *d*=0.19. A sample of at least 870 participants (in a between-subject design), or 435 subjects (in a within-subject design), which is over 26 to 54 (within-subject 13 to 26) times greater than previous investigations, would be required to reliably detect even this “optimistic” negative effect size estimate with statistical power of 0.8. Regression analyses with comprehensive controls corroborate the absence of a main treatment effect, and the absence of moderation by 2D:4D (right hand, left hand, and their average), as implied by insignificant interaction coefficients (see Supplementary Material, Table S3a). Furthermore, in an analysis analogous to the previous positive report [17], we found no correlation between the treatment effect on the RMET and the right-hand 2D:4D in the testosterone group (*r*(123)=0.04, *P*=0.66, 95% CI=[−0.14, 0.22]) (see Fig. S4 in the Supplementary Material).

Experiment 2, which had 400 participants, could also not reject the null hypothesis (beta=0.27, 95% CI=[−0.49, 1.02]; *t*(398)=0.69, *P*=0.49; Cohen’s *d*=0.04, 95% CI=[−0.15, 0.24]) and there was no significant treatment effect in any regression model (Supplementary Material Tables S3b and S4b). Similarly, the point estimate of the effect in Experiment 2 was positive, and the 95% CI did not include negative effects of testosterone administration on the RMET that were greater in magnitude than 0.15. Further analyses of each question in isolation using Chi-squared tests revealed no systematic differences between treatment conditions in any of the RMET items in both experimental datasets (see Supplementary Material Tables S5a and b).

### Testing for effect of 2D:4D

Putative prenatal testosterone proxies (2D:4D, either right-hand, left-hand, or their average) did not correlate with RMET scores in both experiments or moderate the effect of testosterone administration, echoing other recent findings [14–16] (see Supplementary Material Tables S3a-S4b). These results are in line with previous reports showing no correlation between the 2D:4D and RMET scores [12,38,39], and in contrast to the two papers reporting an interaction between 2D:4D and the exogenous testosterone’s effect on the RMET [14,17].

## Discussion

Our experiments used two notably large samples to test the effects of pharmacological testosterone manipulation on cognitive empathy. Despite experimental differences between them, their collected data exhibit the same results with robust statistical consistency, to demonstrate a lack of effects of testosterone administration and 2D:4D on cognitive empathy. These findings, and the literature as a whole, cast serious doubts on the proposal that testosterone causally impairs cognitive empathy, for several reasons.

First, the low statistical power of previous investigations undermines their reliability in capturing true effects. Even if we assume that a purported size of testosterone’s negative effect on cognitive empathy is the overly “optimistic” negative bound of our confidence interval for *d*=−0.19 in Experiment 1, we find that all previous investigations of the topic were statistically underpowered (<0.3 power). Second, the results of the previous small sample studies are discrepant. Our large samples draw on drastically more data than all previous investigations combined, and generalize across geographically, economically, and culturally distinct populations (see Participants section of Supplementary Material). Our use of two different experimental designs and testosterone administration protocols across these populations further mitigates the concern that the outcomes were due to a particular experimental factor. Of note, there are some design differences between our studies and previous investigations (see Table 1; differences from [17] discussed above). However, even if those design differences led to a complete abolishment of a “real” effect of testosterone on cognitive empathy, our results demonstrate beyond a reasonable doubt that such an effect is not generalizable to both males and females. Future work with females could employ a similar approach as ours characterized by large samples from different geographies, distinct administration methods, and other design features that strongly inform whether a relationship (or its absence) generalizes across sexes.

**Table 1.** Summary of literature linking testosterone administration and the RMET

| Study design |  |  |  |  | Results |  |  |  |  |  |
| --- | --- | --- | --- | --- | --- | --- | --- | --- | --- | --- |
|  | Sex | N | Design | Dose | Main effect | Effect Size | SE of Effect Size | 95% CI for Effect Size |  | 2D:4D moderation |
| van Honk et al. (2011) | F | 16 | Within subject; repeated task | 0.5 mg sublingual | One-tailed Wilcoxon<br>P = 0.01 | -0.49 | 0.25 | -1.19 | 0.21 | Low right hand ratio, negative relationship |
| Olsson et al. (2016) | F | 33 | Within subject; repeated task | 50 mg transdermal | One-tailed<br>P = 0.048 | -0.33 | 0.13 | -0.15 | 0.82 | No |
| Bos et al. (2016) | F | 16 | Within subject; no repetition | 0.5 mg sublingual | No effect<br>P = 0.78 | -0.10 | 0.10 | -0.60 | 0.79 | No |
| Carré et al. (2015) | M | 30 | Within subject | 150 mg transdermal | No effect<br>P = 0.25 | 0.20 | 0.35 | -0.70 | 0.31 | Low left hand ratio, negative relationship |
| Nadler et al. Experiment 1 | M | 241 | Between subject; no repetition | 100 mg transdermal | No effect<br>P = 0.83 | 0.03 | 0.36 | -0.19 | 0.28 | No |
| Nadler et al. Experiment 2 | M | 400 | Within subject; no repetition | 11 mg intranasal | No effect<br>P = 0.66 | 0.04 | 0.26 | -0.15 | 0.24 | No |

In "repeated task" paradigms, participants completed the RMET twice: prior to any treatment in Experiment 1 in this study, and after receiving a placebo or testosterone in van Honk et al. (2011).
Effect size is calculated using Hedge's bias-correct effect size due to small sample size and unequal variances between treatment and control groups.

A third reason concerns the validity of the 2D:4D biomarker. The initial findings that prenatal testosterone exposure correlates with 2D:4D are supported in non-clinical and clinical human populations [11], as well as in preliminary causal evidence in relative phalanx/tibia lengths in mice [18]. However, recent work highlights concerns regarding the reliability of 2D:4D as a biomarker [41,42]. For example, the 2D:4D of complete androgen insensitivity syndrome patients were found to be only somewhat feminized, and had the same variance as in healthy controls, demonstrating that the preponderance of individual differences in the measure is not attributable to the influence of testosterone exposure [19]. There is also longitudinal evidence that 2D:4D systematically changes during childhood [43,44], which is unconformable with the preposition that it accurately quantifies prenatal influences. Moreover, while many studies report 2D:4D sexual dimorphism [45,46], other studies suggest lack of ethnic universality of dimorphism [47,48]. Finally, there is also a debate on whether sexual dimorphism is the product of allometric shift in shape rather than hormonal influences [49,50].

Furthermore, many reports of correlations between 2D:4D and behavioural traits hold only for subsets of the population (e.g., particular sex or race). Correlation sometimes holds only for the right hand 2D:4D but in other times only for the left hand, or for the average of both hands [51]. Overall, significant results are seldom replicated, and few survive correction for multiple comparisons or meta-analytic aggregations (e.g., [52]). These concerns belie the validity of the measure as a biomarker and its capacity to detect reliable correlations with noisy psychological constructs in studies of small samples.

Despite our dissenting results, the absence of evidence is not necessarily evidence of absence. Specifically, the lack of an association between 2D:4D and cognitive empathy could be attributable to the failure of the measure to serve as a reliable androgenic biomarker. We therefore agree with Baron-Cohen et al. [53] (p. 6) that it is worthwhile to study the occurrence of impaired cognitive empathy and other ASD traits in developmentally unique populations. One such study reported mixed evidence of higher scores in some Autism Quotient self-rating subscales among women with congenital adrenal hyperlexia (CAH) and lower scores in other subscales, compared with their unaffected relatives [54], with no significant results in men. However, results along this line of research, too, are far from being conclusive. For example, [9] found that young females with and without CAH did not differ in autistic traits, and that amniotic testosterone levels were not associated with scores from either sex individually or the entire sample. Other longitudinal studies also found no association between various measures of prenatal androgens measured in umbilical cord blood and amniotic fluid and autistic traits [55,56]. Thus, further investigations, preferably using larger samples, are required for resolving the inconsistencies in this literature.

To conclude, we tested testosterone’s causal role in cognitive empathy across distinct administration methods using notably large samples from two distinct populations, and found no evidence of an effect of testosterone administration on RMET in young adult neurotypical human males. While our results do not exclude all possible relationships between testosterone and interpreting others’ emotions and states of mind, our large-scale study and evaluation of previous literature exhibit robust evidence of no causal relationship between activational and purported developmental testosterone exposure and cognitive empathy.

## Author contributions

A.N.: experimental design, manuscript, data analysis; G.N.: experimental design, manuscript, data analysis; C.C.: manuscript; D.Z.: manuscript, hormonal assay; T.L.O.: experimental design, data collection; hormonal assay; N.V.W.: manuscript; J.M.C.: experimental design, manuscript.

## Acknowledgements

Funding for this work was generously provided by Caltech, Ivey Business School, IFREE, Russell Sage Foundation, University of Southern California, INSEAD, Stockholm School of Economics, Wharton Neuroscience Initiative, the Natural Sciences and Engineering Council of Canada, and the Northern Ontario Heritage Fund Corporation. Special thank you to Jorge Barraza, Austin Henderson, Garrett Thoelen, Dylan Mandred, Kimberly Gilbert, Caelan Mathers, Emily Jeanneault, Nicole Marley, Kendra Maracle, Victoria Bass-Parcher, Nadia Desrosiers, Charlotte Miller, Brittney Robinson, Dalton Rogers, Megan Phillips, Brandon Reimer, Camille Gray, Christine Jessamine, and Brandon Reimer who assisted this study, and David Kimball for LC-MS/MS assay testing.

## Footnotes

1 Cognitive empathy is the ability to interpret others’ emotions and understand their behaviour vis-a-vis their emotional state; this is distinct from emotional empathy, which is the vicarious feeling of others’ emotions along with them (Smith 2006).

2 The DSM V criteria for ASDs include “Non-verbal communication problems, such as abnormal eye contact, posture, facial expressions, tone of voice and gestures, as well as an inability to understand these.”

3 ASD incidence rates vary widely by study, from 5.2 to 72.6 per 10,000 people and ratios range from 1.81 to 15.7 male:female.

4 The primary publication [17] had a statistical power of only 0.26 to detect the effect size found in the similar study with twice the sample size [16].

5 One experiment (in males) reported a statistically significant moderating effect, but only for the left-hand 2D:4D; the two other experiments reported no moderation of the 2D:4D [15].

6 The RMET has a test-retest reliability of 0.7 [6].

7 Subsequent studies measuring testosterone in serum in significantly larger sample sizes demonstrate an earlier hormonal peak at 60 minutes post administration with subsequent stabilization [14,25].

8 Median testosterone levels (unlogged) of the testosterone group were 33.5 times that of the placebo group post-treatment.

9 These include RMET baseline scores, portion A or B, 2D:4D and treatment interactions, Cognitive Reflection Task (CRT) scores, math abilities, mood and affective measures, treatment expectancy, age, marital status, sexual preference, and all other measured hormones that were not influenced by testosterone treatment and may affect cognition and decision-making (e.g., cortisol [35,36]). The CRT control was added because performance is impaired by exogenous testosterone [29] and people with ASD outperform non-ASD age-matched controls [37].

10 These include Cognitive Reflection Task (CRT) scores, factor 1 and 2 psychopathy measures, treatment expectancy, age, marital status, sexual preference, and all other measured hormones that were not influenced by testosterone treatment.

