## Supplementary material for "Does testosterone impair men’s cognitive empathy? Evidence from two large-scale randomized controlled trials"

**Affiliations:**
[a] Department of Finance, Ivey Business School at Western University, 1255 Western Rd, London, ON, Canada N6G 0N1;
[b] Marketing Department, The Wharton School of the University of Pennsylvania, 3730 Walnut St, Philadelphia, PA, USA 19104;
[c] ZRT Laboratory, 8605 SW Creekside Place, Beaverton, OR, USA 97008;
[d] Department of Psychology, Nipissing University, 100 College Drive, North Bay, ON, Canada, P1B 8L7;
[e] Department of Psychology, Simon Fraser University, 8888 University Drive, Burnaby, BC, Canada, V5A 1S6;
[f] Department of Psychology, Nipissing University, 100 College Drive, North Bay, ON, Canada, P1B 8L7;
[f] Department of the Humanities and Social Sciences, California Institute of Technology, 1200 E California Blvd, MC 228-77, Pasadena, CA, USA 91125;

**This PDF file includes:**

Supplementary text
Figs. S1a to S4
Tables S1a to S7b
References for Supplementary Materials reference citations

**Other supplementary materials for this manuscript include the following:**

Datasets S1 and S2

### Participants

**Experiment 1.** Most participants (217, 90%) were students from The Claremont Colleges, a private southern California college consortium, or nearby post-secondary institutions; non-student participants were community members from surrounding cities. Of an initial sample of 244 male participants, two did not complete the task and one did not show up for the afternoon session; those three were excluded from further analysis (final *N* = 241). *N* = 123 were assigned to receive a standard dose of 100mg transdermal testosterone (drug name is Vogelxo™) and *N* = 118 to receive placebos, in a pre-randomized, placebo-controlled double blind exogenous administration design (see Fig. S1a). Each session dispensed either testosterone or placebo such that there were no sessions with "mixed" testosterone and placebo treatments provided. Participants' ethnic backgrounds were diverse and consisted of 45% White/Caucasian, 22% Black, 10% South Asian, 5% Latin American, 5% Asian, 3% South East Asian, 2% Arab/West Asian, and 8% reported "other & mixed".

Pre-screening criteria excluded participants with relevant medical and psychological conditions (i.e., 5α-reductase deficiency, Klinefelter syndrome, brain tumor, cancer, psychiatric diagnosis/diagnoses, high blood pressure, liver disease, kidney disease, angina, cancer, hepatitis, renal/kidney impairment, history of epileptic seizures, and hypersensitivity to soy/alcohol), those using prescription drugs that could interfere with the study (i.e., oxyphenbutazone, insulin, corticosteroids, and opioids), those who self-reported having consumed illegal drugs or excessive alcohol in the previous 24 hours, and non-fluent English speakers.

Personal, demographic, and treatment expectancy characteristics of all participants in the two treatment groups are summarized in Table S1a (note that five participants did not report their age and were therefore excluded from all analyses in which age is used as a control variable). The right column of Table S1a also reports the *P*-values of two sample *t*-tests that assessed differences in characteristics between the testosterone and placebo groups (a check on whether random assignment resulted in balance on all such variables). Two participants (one from each treatment group) self-reported taking testosterone treatment on a regular basis; all analyses include these participants and are robust to excluding them. To reduce the potential effect of a female researcher's presence on testosterone-related behaviors, exclusively male researchers conducted all experimental sessions.

Since all sessions were exclusively testosterone or placebo, we aimed to achieve a ratio of 1 for testosterone/placebo administration. Given differences in show-up between sessions, there was a known possibility that there would be an imperfect ratio, although it would be practically and statistically immaterial. See Figure S1a for the flow diagram depicting the phases of the recruitment, eligibility, and randomization processes.

**Experiment 2.** The sample of 400 participants, 130 from Sudbury and greater area in northern Ontario, Canada, and 270 from North Bay and surrounding area, was comprised of both students and general public. All 400 accepted male participants completed the task and were included in the analysis. Using a predetermined random sequence (generated by www.random.org), 199 participants were assigned to receive 11 mg intransal testosterone (Natesto®) and 201 participants were randomly assigned to receive placebo, in a randomized, placebo-controlled double-blind exogenous administration design.

These final samples exclude 32 men who failed the intake process for any disqualifying condition (e.g., heart condition, diagnosed with psychological or developmental disorder, drug or alcohol dependence, were on sport team for which testosterone is a banned substance). See Figure S1b for the flow diagram depicting the phases of the recruitment, eligibility, and randomization processes.
Upon registration for the study, participants were excluded from participating if they were not between the ages of 18 and 40, on a sports team where testosterone is a banned substance, drug or alcohol dependent, on any prescription medication known to interfere with steroid hormone concentrations (e.g., corticosteroids, anabolic steroids), have a medical condition such as prostate cancer, diabetes, or heart condition, or be currently diagnosed with any developmental or psychological disorders (e.g., schizophrenia, bipolar disorder, borderline personality disorder, anxiety disorder, depression PTSD, obsessive compulsive disorder, attention deficit hyperactive disorder, panic disorder).

Personal, demographic, and treatment expectancy characteristics of all participants in the two treatment groups are summarized in Table S1b. The right column also reports the *P*-values of two sample *t*-tests that assessed differences in characteristics between the testosterone and placebo groups (a check on whether random assignment resulted in balance on all such variables). Participant ethnicities were as follows: 80% White/Caucasian, 7% Native, 6% Black, and 7%

reported “other & mixed”.

We aimed to achieve a ratio of 1 for testosterone/placebo administration. One participant who received testosterone dropped out of the study shortly after drug administration, so to maintain our target of $N = 400$ men, we replaced him with an additional participant who, by chance, received placebo. This caused the slightly unbalanced number of people who received testosterone ($N = 199$) and placebo ($N = 201$).

### Experimental Procedures

**Experiment 1.** Participants first arrived at the lab at 9:00 a.m. for their experimental session. They signed an informed consent form, received a private participant number to ensure anonymity, and then proceeded to a designated room where their hands were scanned (to obtain digit ratio measurement, a putative proxy of prenatal testosterone). Participants were then randomly assigned to private cubicles where they completed demographic and mood questionnaires, and the first portion of the Reading the Mind in the Eyes Test (RMET; see Figure S2 for example item), a widely used 36-item multiple-choice task that measures participants' ability to infer people's mental states by having them choose which of four words best describes what a person is thinking or feeling based on an image of their eye region. The participants were under no time pressure and received no performance feedback.

After all participants completed those questionnaires, they provided an initial baseline saliva sample. Participants then proceeded to the gel administration room for testosterone or placebo gel application (for more details, see *Treatment Administration* below). In the process of signing consent forms, participants were explicitly told to have no skin contact with females, to avoid vigorous exercise and showering, to finish eating their lunch no later than 1:00 p.m., and to return to the lab at 1:55 p.m. well hydrated. These instructions and precautions were repeated again before they departed from the morning session.

All participants returned to the lab prior to 2:00 p.m. (with no incidents of lateness). They provided a second saliva sample and proceeded to the behavioral experiment. The time frame between gel application and behavioral experiment was chosen so that tasks took place when the testosterone group participants experienced elevated and stable blood testosterone levels following drug administration (Eisenegger et al. 2013) (see manuscript Figure 1).

The full-day session consisted of seven independent behavioral tasks measuring cognitive reflection (Frederick 2005; Nave et al. 2017), risk preferences (Holt and Laury 2005; Gneezy and Potters 1997), math ability, brand preferences (Nave et al. 2018), competitiveness (Niederle and Vesterlund 2005), and a die roll task (Henderson et al. 2018). None of the tasks included feedback about the participants' monetary payoffs or relative standing prior to other tasks to minimize changes in testosterone from changes in payoff or winning or losing (only the final task provided feedback regarding each participant's performance relative to other participants in the session). The second portion of the RMET was the second task in the battery, and it took place at approximately 2:20 p.m. (20 minutes following the first post-treatment saliva sample).

The rationale for conducting a battery of tasks (instead of a single experiment) is to maximize the knowledge gained from each human subject undergoing a pharmacological manipulation, which is a standard practice (Zethraeus et al. 2009; Kocoska-Maras et al. 2011) that is viewed favorably by an institutional review board (IRB). This study was approved by Claremont Graduate University's IRB (#2295). No harms or adverse events occurred in this experiment.

To maintain high-resolution monitoring of hormonal changes and control for their influences, a total of four saliva samples were collected throughout the experiment (for details of collection frequency and time, see *Pre-and Post-Administration Saliva Samples* below). The accuracy and consistency of sampling times are crucial because of the measured hormones' unique diurnal cycles, which complicate comparing samples taken at different times of day. To standardize hormonal measurements among all participants, we did not randomize the order of the behavioral tasks, in a similar fashion to previous studies (Zethraeus et al. 2009; Kocoska-Maras et al. 2011). The behavioral battery of tests lasted approximately two hours. The behavioral task reported here occurred approximately 20 minutes after the second saliva sample. Following the experiment, participants completed an exit survey and indicated their guess regarding which of the two treatments they had received (see Table S1a). A research assistant paid participants privately based on their unique identification number at the conclusion of each session.

Participants were tested in 17 distinct cohorts of 13-16 participants at a time with two research assistants staffing each session. Data collection took place over just under two-and-a-half months (September 10$^{th}$, 2014 to November 22$^{nd}$, 2014).

**Experiment 2.** Participants arrived at one of three testing session times, 10:00 a.m., 12:30 p.m., or 2:30 p.m. Each participant was brought into a private testing room upon arrival, and rooms were assigned based on alphabetical order of the room names and first come first served for the participants. When settled into their testing room, participants read and signed an informed consent form and received a participant number to ensure anonymity. Upon signature, the research assistants asked the participants to turn off any devices they had brought with them, to minimize any distractions throughout the study. The first component of the study was a battery of self-report questionnaires averaging 20-30 minutes in duration. It was at this time that participants completed demographic questionnaires, the Self-Report Psychopathy Scale, and other personality-based measures used as part of a larger protocol. After completing the questionnaires, participants then provided a 1-2 ml saliva sample to measure baseline testosterone, and a 50 ml mouthwash sample to be used for genetic analysis (e.g., androgen receptor polymorphism) for a separate research study. Once both samples were collected, a research assistant demonstrated the administration protocol for the drug while reading from a script (see *Treatment Administration* below).

Treatment administration was followed by a 30-minute wait period, during which participants had facial photos taken and their hands scanned (once the 30 minutes had passed, participants had been in the lab for approximately one hour at this point). They then completed three computerized tasks (Point Subtraction Aggression Paradigm, Public Goods Game, and Cognitive Reflection Test) before completing the RMET task. All tasks were performed in the same order and participants were informed of how much their partner contributed in the Public Goods Game. Participants evaluated all 36 items of the RMET task at this time and provided their final saliva sample shortly after. The second and last saliva sample was taken at the completion of all tasks, which fell approximately two hours after arrival to the lab and one-and-a-half hours after drug administration. If participants completed the tasks faster than others, they were asked to wait until the two-hour mark of their study completion to keep sampling consistent across participants. At the completion of the study, participants received compensation ($20 an hour with up to an additional $5 to $20 for their performance on varying tasks), signed a receipt, and completed an exit survey asking which treatment they believed they had received (see manuscript Figure 1).

Participants were tested in cohorts of six at a time with three research assistants each monitoring two participants. Participants were tested in individual testing rooms. Data collection took place over 30 days of sessions taking place between November 28th, 2016 and April 27th, 2017. This study used both male and female research assistants. Results reported in this manuscript are robust to controlling for research assistant sex (see Table S3b). The Nipissing University Research Ethics Board approved this study (REB #100735). No harms or adverse events occurred in this experiment.

#### Treatment Administration Procedures

**Experiment 1.** After providing the first saliva sample, participants were escorted in groups of two to six to a semi-private room where a research assistant provided a small plastic cup containing clear gel and stated it was equally likely to contain testosterone or placebo. To ensure the treatment was double blind for both assistant and participant, the lab manager filled the cups in advance, did not interact with participants, and did not reveal the contents of the cup to the research assistant. These cups contained either 10 g of topical testosterone 1% (two packets of Vogelxo™ by Upsher-Smith, each delivering 50 mg of testosterone in 5 g of gel for a total of 100 mg of T) or the volume equivalent of an inert placebo of similar texture and viscosity (80% alcogel, 20% Versagel®). Based on demonstrated inability to detect treatment through two survey questions, testosterone and placebo were not differentiable by participants (Table S1a).

We chose to administer testosterone using topical gel in Experiment 1 because the pharmacokinetics of a single-dose administration (i.e., a time-course of post-treatment testosterone levels change) have been investigated in healthy young men (Eisenegger et al. 2013) for this specific testosterone administration method. The single-dose study demonstrated that plasma testosterone levels peaked three hours following exogenous topical administration, and that testosterone measurements stabilized on high levels during the time window between four and seven hours following administration. Therefore, all participants were instructed to return to the lab four-and-a-half hours after receiving the gel, when their androgen levels would be higher and more stable. A newer study (Carré et al. 2015) showed a spike in serum testosterone occurring at 60 min relative to 120 min, further supporting our research design of having participants return to the lab at after a possible initial spike and at stable levels.

Participants were instructed as follows:

*"Welcome to the goop room! Have a seat in any chair you want.*

*In this container there is either testosterone or placebo; you have a 50-50 shot. Either way the gel is a lot like hand sanitizer—it will evaporate quickly, so you just need to wait for it to dry before putting your shirt back on.*

*I'm going to go over a few points, shirt off, here's how to apply (demonstrate scooping it out, rubbing on shoulders and upper trapezius). I talk; you apply.*
*You are a healthy male and there are no real hazards, but there is a precaution with testosterone in regards to women. Avoid physical contact with females today; keep your shirt on if you do come in contact. Reason for this is the possibility of serious harm if she's pregnant.*

*No showering or vigorous exercise until after the second part of the experiment.*

*Make sure to finish eating lunch no later than 1:00pm and be hydrated when you come in at 1:50pm today. Please note: if you are late, all of the other participants will have to wait for you before we start the experiment.*

*There is no deception at this experiment or any other at this lab, everything is at is seems. Wash hands as soon you walk out the door before you touch anything, there's a bathroom to the left as well as a sink in the kitchen; head back to the computer lab OR get checked out at the door before you head out. Please read the list of precautions carefully (Hand them Morning Process Form). It is extremely important that you comply with the instructions completely."*

Participants removed upper-body clothing and applied the entire contents of the gel container to their shoulders, upper arms, and chest, as demonstrated by gestures made by the research assistant while reading the above scrip. After self-administering the gel under the supervision of the experimenter, participants were instructed as per medial precautions prior to dismissal as recommended by the gel manufacturers.

All surfaces in the administration room were covered with medical-grade isolation sheets, and surfaces in the gel application area were cleaned with alcohol swabs after each experimental session. The adjacent bathroom where the sink was located was also thoroughly wiped, as were doorknobs and handles.

**Experiment 2.** Once the initial saliva sample was collected, a researcher entered the participant's private testing room with two syringes each containing 5.5 mg of either placebo or testosterone gel. The syringes had been filled in advance by a pharmacist, who did not interact with participants and did not reveal the contents of the syringes to the research assistant performing the study so that the treatment was double blind for both research assistant and participant. A recent study indicates that serum testosterone concentrations rise sharply within 15 minutes after Natesto® gel application and remain elevated for approximately three hours post application among hypogonadal males (Geniole SN, Procyhyn TL, Marley N, Or...) (see Fig. S3). Random assignment was determined such that half the participants in every group received testosterone and half received placebo. Based on demonstrated inability to detect treatment through survey questions, testosterone and placebo were not differentiable by participants (Table S1b).

For the drug application instructions, research assistants had to read and/or memorize the following script for the drug administration.

*"Now we are moving onto the drug administration portion. Remember this is a double blind study, meaning I do not know what drug you are receiving and neither do you. You have a 50% chance of receiving either testosterone or placebo. Don't worry, these are not needles, they are just syringes. The gel inside is very similar to Vaseline and doesn't need to go too deep into your nose."*

The research assistant then read the following administration script:

*"Now you are going to administer the gel. Here are the syringes, you are going to place your fingers on the white label and place the syringe in your left nostril first till your fingers touch your nose.*
*Make sure the syringe is angled to the outer edge of your nose. Once at the proper location, place your other hand on the applicator and slowly empty the syringe while scraping it along the outer edge of your nose. Understand?*
*When you're done, place the syringe in the garbage and then we will do the other side. Ready? There's a mirror here if you need it. Now we are going to do the other side. Now pinch your nose with both your fingers to make sure the gel has spread all around the inside of each nostril. Here is some water in case any of the gel seeps down into your mouth. This way you can wash it down. Please refrain from sniffling, touching, or blowing your nose.*

*Now we have a 30-minute wait period before we can do any additional testing. There will be a couple of tasks in the meantime, so just sit tight and I will come in and out with the tasks."*

Once the administration of the gel was complete, the research assistant pointed to the box of tissues and hand sanitizer and instructed the participants to sanitize thoroughly before touching any surfaces in case any there was any contamination of the drug on the outside of the syringes.

**Pre- and Post-Administration Saliva Samples**

**Experiment 1.** Each participant provided two saliva samples at predetermined sampling times: (1) before treatment administration (all samples provided between 9:25 and 9:34 a.m., 9:27 a.m. on average, *SD* = 3.4 minutes), and (2) upon return to the lab, just prior to starting the behavioral tasks (all samples provided between 1:55 and 2:15 p.m.; 2:02 p.m. on average, *SD* = 5.3 minutes). Two additional saliva samples were taken later in the experiment to quantify hormonal changes occurring in other parts of the experiment. We chose to assay saliva samples rather than serum to avoid stress associated with repeat venipuncture and because saliva serves as a reliable measure of exogenous testosterone (Mayo et al. 2004; Du et al. 2013). Time-stamped saliva samples were acquired by participants by collecting passive drool into plastic containers at their private workstations, as instructed by research assistant (see manuscript Figure 1).

Fourteen hormone measures were acquired using liquid chromatography tandem mass spectrometry (LC-MS/MS) (see Table S3a) conducted by ZRT Laboratories. No food or drinks were allowed into the laboratory.

**Experiment 2.** Each participant provided two saliva samples, one before drug administration, and one at the completion of the study. Times varied depending on testing session: samples were collected at approximately 10:30 a.m. and noon (10:00 a.m. session), 1:00 p.m. and 2:30 p.m. (12:30 p.m. session), or 3:00 p.m. and 4:30 p.m. (2:30 p.m. session). Participants provided passive drool into a 5 ml polystyrene tube in their individual testing rooms as instructed by a research assistant. All saliva samples were analyzed using commercially available enzyme immunoassay kits (DRG International) (see Table S2b).

**Hormonal Assay Procedures**

**Experiment 1.** Salivary steroids (i.e., estrone, estradiol, estriol, testosterone, androstenedione, DHEA, 5-alpha DHT, progesterone, 17OH-progesterone, 11-deoxycortisol, cortisol, cortisone, and corticosterone) were measured by LC-MS/MS using an AB Sciex Triple Quad 5500. Internal standards were added to 1 ml of saliva, and the steroids then extracted by C18 column chromatography with 0.1 M $NH_4OH$ wash followed by 10% acetone. Steroids were eluted from the SPE with 10% methanol in acetone and dried under nitrogen. The dried samples were subjected to derivatization—the process of transforming a compound into a derivative product of similar chemical structure—with pyridine-3-sulfonyl chloride for the estrogens (i.e., estrone (E1), estradiol (E2), and estriol (E3)), as outlined by (Xu and Spink 2008). Added to the dried samples were 40 µL sodium bicarbonate (50 mM, pH 10) and 40 µL pyridine-3-sulfonyl chloride (3 mg/mL in acetonitrile). The resulting mixture was then incubated at 60C° for 10 min. After derivatization, the samples were diluted with 80 µL of water and injected for LC-MS/MS analysis with analytical separation performed on an Agilent Poroshell 120 EC-C8 column and ionization by atmospheric pressure chemical ionization (APCI) in the positive ionization mode. Table S8a lists each analyte and its validation results for the lower limit of quantitation (LLOQ, which is jargon for the lowest level of detection with coefficients of variation (CVs) < 20% over the linear range), linear range, and the inter-assay precision from the highest concentration to the LLOQ within the linear range. When salivary hormone levels of participants were below their LLOQ, we assigned values halfway between zero and their respective LLOQ (note that the true quantities of the hormone in the sample are never zero, even when they do not reach the detection threshold).

**Experiment 2.** Salivary steroids (i.e., testosterone and cortisol) were measured using commercial enzyme immunoassay kits (DRG International). All samples were assayed in duplicate and the average of the duplicate samples were used for statistical analyses. The intra- and inter-assay CVs were below 10%. Table S8b shows detection levels, precision, and normality tests.

All hormone data (on both studies) were normalized via log transformation prior to statistical analyses (for Kolmogorov-Smirnov normality tests of the non-logged and logged hormonal measurements; Tables S2a and S2b present raw values for clarity).

**Hormonal Changes Following Treatment and Manipulation Check**

**Experiment 1.** As expected, we found significant post-treatment differences between groups with respect to all hormones influenced by testosterone treatment, either as an upstream (androstenedione) or downstream (5-alpha DHT) metabolite of testosterone (Horton and Tait 1966): logged post-treatment testosterone group mean testosterone levels 8.38 = pg/ml $SD = 0.15$; logged placebo group mean testosterone levels = 5.11 pg/ml, $SD = 0.08$; two-sided t-test: $P < 0.0001$, $t(239) = 18.43$, and statistically similar pre-treatment testosterone levels: logged pre-treatment testosterone group mean testosterone levels = 5.58 pg/ml, $SD = 0.08$; logged Placebo group mean testosterone levels = 5.77 pg/ml, $SD = 0.09$; two-sided t-test: $P = 0.13$, $t(239) = 1.50$. Two additional samples were taken later during this day-long experiment to provide measures for other tasks: sample C testosterone levels in the testosterone group were 9.23, SD = 1.45 and the placebo group mean was 5.28, SD = 0.962; two-sided t-test: P < 0.0001, t(240) = 24.8. The fourth samples were similar, with the testosterone group mean testosterone levels of 9.16, SD = 0.13, placebo mean testosterone levels of 5.19, SD = 0.92; two-sided t-test: P < 0.0001, t(240) = 25.8. Additionally, we found a decrease in progesterone 17OH resulting from an increase in testosterone, which is commonly observed in commercial laboratory assays following testosterone supplementation. The changes in saliva testosterone measures were similar in magnitude to those reported in previous studies following topical gel administration of testosterone and progesterone (Hurwitz, Cohen, and Williams 2004; Hucklebridge et al. 2005) (see manuscript Figure 1, Table S2a and dataset S1).

We observed no significant differences between treatment groups in the hormones that were not expected to change following short-term testosterone treatment (e.g., aldosterone, cortisol, melatonin) in pre-treatment and post-treatment saliva measurements. The pre-treatment and first post-treatment mean hormonal saliva levels are summarized in Table S2a; note that differences between morning and afternoon hormonal levels were affected by diurnal cycles in both treatment groups.

From assays conducted during the first 13 (out of 17) sessions of the study, we noted that 72 out of 184 pre-treatment baseline saliva samples (in both treatment groups) presented measurements with higher testosterone levels that are expected in normal young men (greater than 400 pg/mL). All other hormone measurements (including testosterone metabolites) were typical. We traced the cause of these abnormal measurements to testosterone gel transfer to common surfaces (e.g., doorknobs and mouse pads). Crucially, the high measurements were caused by the local spread of testosterone into saliva tubes, but physiological levels were unaffected by superficial contact with the dry nuisance testosterone gel, as (a) we observed normal pre-treatment levels of testosterone metabolites, namely DHT and androstenedione in all subjects; (b) none of the placebo group participants showed abnormally high values of testosterone metabolites in any of the post-treatment measurements; (c) only five out of 118 subjects from the placebo group showed consistently elevated testosterone measurements in all of the three post-treatment saliva samples; and (d) previous investigations found that interpersonal testosterone transfer is highly unlikely even with skin-to-skin contact (Rolf et al. 2002). Thus, we found convergent evidence that biofluid levels were unaffected by superficial contact. This conclusion was supported by evidence found in commercial labs.

In response to this finding, during the course of the experimental period, we identified all surfaces and objects through which testosterone could spread in the facility, and improved the sterile isolation protocol to reduce the spread of the dried testosterone gel. This protocol included using a bleach-alcohol solution to thoroughly clean keyboards, computer mice, chair backs, displays, and all doorknobs after each session and asking subjects to carefully wipe their hands with a wet tissue before collecting each saliva sample. New pens were used for each session, and all previously used pens were removed from the testing area. Clipboards and other miscellaneous objects that participants did, or could, interact with were cleaned, and an aerosol "air sanitizer" that bonds to VOCs (volatile organic compounds) was sprayed into the air. Following the adoption of this strict sterilization protocol, we found a drastic reduction in incidence of high testosterone samples in the pre-treatment measurements, to a total of five participants out of 58 in the following four sessions (sessions 14–17). Each regression analysis was replicated for robustness with exclusion of abnormal testosterone samples and results are virtually unchanged (see *Robustness checks* below).

**Experiment 2.** As expected, we found significant post-treatment differences between groups with respect to testosterone between treatment conditions: logged post-treatment testosterone group mean testosterone levels = 8.00 pg/ml *SD* = 1.27; logged placebo group mean testosterone levels = 5.33 pg/ml, *SD* = 0.73; two-sided t-test: $P < 0.001$, $t(396) = 22.14$. There were no significant differences in baseline testosterone concentrations in pre-treatment testosterone levels: logged testosterone group mean testosterone levels = 5.3 pg/ml, *SD* = 0.88; logged Placebo group mean testosterone levels = 5.4 pg/ml, *SD* = 0.85; two-sided t-test: $P = 0.49$, $t(394) = 0.69$ (See manuscript Figure 1 and Table S2b). Also, there were no differences in cortisol measures between treatment groups: testosterone group mean logged cortisol = 0.95 ng/ml, *SD* = 0.71; placebo group logged cortisol meaN = 0.91 ng/ml, *SD* = 0.71; two-sided t-test: $P = 0.51$, $t(395) = 0.66$ (see dataset S2).

**Digit Ratio Measurements**

The ratio of second (index) finger length to fourth (ring) finger (abbreviated 2D:4D) is considered a proxy for prenatal testosterone exposure (Manning 2011), although this hypothesis is still under debate (e.g., (Valla and Ceci 2011)).

**Experiment 1.** Participants' 2D:4D were measured by two independent raters using hand scans and digital calipers. Correlation between the two raters was 0.95 and if measures differed by over 5%, the participants were flagged and remeasured. The right-hand digit ratio was not calculated for one subject due to a broken finger, and he was therefore excluded from all analyses that used the right-hand digit ratio as a control.

Correlation between the digit ratios of the left and right hands was 0.64, $P < 0.0001$ (Table S6a). The two treatment groups did not differ with respect to left-hand 2D:4D ($t(240) = 0.57$, $P = 0.57$), and the average of two hands ($t(239) = 1.48$, $P = 0.14$). A slightly lower (i.e., more "masculinized") right-hand 2D:4D was identified in the testosterone group (T group meaN =

0.943, placebo group mean = 0.953, $t(240) = 2.27$, $P = 0.024$—corrected P-value threshold should be 0.05/3 = 0.017, which is not met). Digit ratio had no impact on RMET scores as shown by regression analysis.

**Experiment 2.** Participants' 2D:4D were measured by two independent raters using hand scans and digital calipers (correlation between the two raters was 0.86). A scanner malfunction caused loss of data for the first 33 participants.

Correlation between the digit ratios of the left and right hands was 0.59, $P < 0.0001$ (Table S6b). The two treatment groups did not differ with respect to left-hand 2D:4D (T group mean = 0.945, $SD = 0.03$ placebo group mean = 0.948, $SD = 0.03$ $t(354) = 0.79$, $P = 0.43$), right hand 2D:4D (T group mean = 0.951, $SD = 0.03$, placebo group mean = 0.953, $SD = 0.03$; $t(355) = 0.98$, $P = 0.33$) or the average of two hands (T group mean = 0.948, SD = 0.03 placebo group mean = 0.951, $SD = 0.03$; ($t(354) = 0.965$, $P = 0.34$) (Table S6b). Similar to Experiment 1, digit ratio had no impact on RMET scores as shown by regression analysis.

**Psychological Questionnaires**

**Experiment 1.** We measured mood using the PANAS-X scale (Watson and Clark 1999), both pre-treatment (in the morning) and post-treatment (in the afternoon). Table S1a shows no significant difference between treatment groups. We found a modest decrease in both affect measures over time (morning vs. afternoon), and no treatment or time × treatment interaction, indicated by the output of two-way analysis of variance (ANOVA) with an interaction term, ruling out this indirect way in which testosterone might affect social cognition or behavior. Three participants did not answer all of the negative-affect items in their questionnaires, and five participants did not complete all of the positive-affect items; these participants were excluded from analyses that included these scales as control variables.

**Experiment 2.** A previous study proposed that psychopathic traits moderate the effect of testosterone administration on the RMET (Carré et al. 2015). We measured psychopathic traits using the Self-Report Psychopathy-Short Form (SRP-SF; (Paulhus and Williams 2002)). This questionnaire consists of 29 items covering four factors: Interpersonal, affective, lifestyle and antisocial which collapse into two factors. Factor 1 consists of the interpersonal and affective items, and Factor 2 consists of the lifestyle and antisocial items. Each question was presented with five possible responses on a Likert scale (strongly disagree to strongly agree). We found no difference between treatment groups pre-application ($P$s > 0.78) (see Table S1b).

**Treatment Expectancy**

One previous study suggested an effect of participants' beliefs about the treatment they had received on behavior (Eisenegger et al. 2010).

**Experiment 1.** We asked participants to use a five-point scale to indicate their expectancy about whether they had received placebo or T. Two participants did not report their treatment expectancy and therefore were excluded from all analyses in which this measure was used as a control. There were no significant differences between the groups on this expectancy measure (Table S1a).

**Experiment 2.** We asked participants to guess as to whether they received placebo or testosterone (binary response). There were no differences between the groups on this measure (Table S1b).

**The Reading the Mind in the Eyes Test**

**Experimental Task.** We administered the adult version of the RMET developed by Baron-Cohen and colleagues (Baron-Cohen et al. 2001). In both studies we presented participants with the same rectangular images used in the original study, showing solely the eye region of an actor's face, and a list of four words that describe emotional states. Participants were instructed to select the emotional state that best described the person in the image (see Fig. S2).

We used the following instructions for both experiments:

*"For each set of eyes, choose and click which word best describes what the person in the picture is thinking or feeling.*

*You may feel that more than one word is applicable but please choose just one word, the word which you consider to be most suitable.*

*Before making your choice, make sure that you have read all 4 words.*

*You should try to do the task as quickly as possible but you will not be timed.*

*For your convenience, if you really don't know what a word means you can find its definition at the bottom of each question."*

To reduce the likelihood that participants made choices without fully understanding the word meaning, we included a dictionary defining all words, which is a part of the original RMET materials (Appendix B in Baron-Cohen et al. (2001)).

**Comparability of RMET Scores to Previous Research.** The mean RMET score in Baron-Cohen et al.'s (2001) sample of male students (*N* = 53) was 27.3 (out of 36) (*SD* = 3.7), which is comparable with scores reported by other researchers (e.g., (Smeets et al. 2009)).

**Experiment 1.** Our participants' pooled scores (the sums of both portions) were similar, with an overall average of 27.5 (*SD* = 3.9) (mean score for placebo = 27.3, *SD* = 2.7; mean score for testosterone 27.7, *SD* = 3.4).

**Experiment 2.** The average scores were also similar, with the placebo and testosterone groups scores 25.6 (*SD* = 3.9) and 25.8 (*SD* = 3.9), respectively.

Manuscript Figure 2 compares RMET scores cumulative distribution in Experiment 1 testosterone (*N* = 123) and placebo (*N* = 118) and Experiment 2 testosterone (*N* = 199) and placebo (*N* = 201) groups juxtaposed with samples from general population (*N* = 225). Distribution of correct scores (out of 36) in our sample is similar to non-ASD males and females from general and student populations (2001). Note that experiment 1 and experiment 2 used participants sampled from different and distinct populations.

Manuscript Table 1 provides compilation of results of the experimental literature testing the EMB hypothesis through exogenous testosterone manipulation and the 2D:4D and effects on RMET performance.

Figure S4 exhibits an analogous analysis as done in van Honk et al. (2011) to measure the effect of exogenous testosterone in relation to the 2D:4D. A non-significant result ($r(123) = 0.04$, $P = 0.66$) was found.

**Regressions Analyses: Testing the Effects of testosterone administration and its interaction with 2D:4D on test performance on the RMET**

**Experiment 1.** We estimated ordinary least squares (OLS) regression models with afternoon (post-treatment) RMET scores as the dependent variable (DV) (see Table S4a). Independent variables (IVs) were *Treatment*, a binary variable indicating whether the participant received testosterone (treatment = 1) or placebo; *Morning*, the score on the baseline RMET task taken pre-treatment in the morning; *Order*, a binary variable indicating which portion of the test was taken in the afternoon (such that order = 1 if portion A was taken in morning session and B in the afternoon, and order = 0 if vice versa); *DR_right*, *DR_left* and *DR_average*, the 2D:4D of the right hand, left hand, and their average, respectively; and *DrRTx* and *DrLTx*, the interaction terms for the digit ratio of the right hand and left hand, and their treatment. Other independent variables were hormonal measures that were not affected by the treatment, in logged form, treatment expectancy (belief), number of children, relationship status, married or single, sexual preference, cognitive reflection test score (Frederick 2005), math ability, and positive and negative affect measured using the PANAS-X scale (Watson and Clark 1999). We estimated the following regression models:

*1A estimates the influence of exogenous testosterone (Treatment) on RMET while controlling for Morning and Order.*

*1B includes the digit ratio and interaction variable for treatment and digit ratio for the right hand.*

*1C shows the same specification as (A2) but for the left hand.*

*1D shows the same specification as (A2) but with the average of the digit ratios for the right and left hands.*

*1E shows the right-hand digit ratio and afternoon levels of hormones not influenced by exogenous testosterone (logged values), and the treatment expectancy added to A1.*

*1F builds on Model (A2) and adds affective scores (Watson and Clark 1999) and age.*

*1G builds on model A6 and adds cognitive reflection task (CRT) score and mathematical performance, as the former has been shown to be affected by testosterone (Nave et al. 2017)*

*1H adds relationship, family, and sexual orientation control variables to A7.*

*1I tests the influence of cortisol on cognitive empathy and its interaction with treatment using an interaction term specifying the binary treatment variable and continuous logged cortisol levels.*

The treatment effect in Experiment 1 is defined as the difference between the post-treatment RMET and the baseline scores in the testosterone group, after regressing out the effect of RMET portion (A or B) order. The finding is robust to portion effect and holds when replacing the right-hand 2D:4D with either the left hand or the average of both hands. Results showed no effect of testosterone treatment on the post-administration RMET score (afternoon) in any model. Model 1A beta = 0.106, 95% CI = [−0.45, 0.68], $t(237) = 0.37$, $P = 0.711$. The estimated effect size

(Cohen's *d*) was 0.044, CI = [−0.19, 0.28], which direction is opposite in sign to the hypothesis of the original study. We also found no evidence for moderation of the 2D:4D digit ratio. Expectedly, the morning performance was a significant ($P$ < 0.01) predictor for the afternoon RMET predictors, and so was the order in which the two parts of the test were taken ($P$ < 0.01). The age coefficient was also statistically significant at $P$ < 0.05 in model 1F yet with trivial impact on RMET scores.

To ensure that our results were robust to the exclusion and inclusion of participants associated with samples containing additional testosterone (as discussed above in "hormonal changes following treatment and manipulation check" section), we conducted additional robustness checks by examining the effects of testosterone on the RMET when (a) excluding subjects with pre-treatment saliva testosterone of greater than 400 pg/ml from both the testosterone and placebo groups; (b) excluding placebo subjects with post-treatment saliva testosterone (sample B) greater than 400 pg/ml; (c) excluding all subjects in either condition (a) or (b); and (d) repeating the analysis with a more conservative cutoff of 250 pg/ml for both treatment groups in Experiment 1. We found that the effects of testosterone administration on the RMET were virtually unchanged regardless of the exclusion criteria used (range of betas on model A1 above range from 0.11 to 0.24, all $P$-values for treatment binary variable > 0.44).
Two participants, one from each treatment group, self-reported taking exogenous testosterone regularly. Their overall RMET scores were 22 and 24 (out of 36), which are within 95% confidence within the same sample. Results are robust to their exclusion from the sample.

***Experiment 2:*** We estimated several ordinary least squares (OLS) regression models with RMET scores as the dependent variable (DV) (see Table S4b). Independent variables (IVs) were *Treatment*, a binary variable indicating whether the participant received testosterone (treatment = 1) or placebo; *DR_right*, *DR_left* and *DR_average*, the 2D:4D of the right hand, left hand, and their average, respectively; and *DrRTx* and *DrLTx*, the interaction terms for the digit ratio of the right hand and left hand, and their treatment. Other independent variables were treatment expectancy (belief), number of children, relationship status, married or single, sex of the research assistant (female = 1), and Factor 1 and Factor 2 psychopathy scores.

We estimated the following regression models:

*Model 2A estimates the influence of exogenous testosterone (Treatment) on RMET.*

*2B includes the digit ratio and interaction variable for treatment and digit ratio for the right hand.*

*2C shows the same specification as (A2) but for the left hand.*

*2D shows the same specification as (A2) but with the average of the digit ratios for the right and left hands.*

*2E shows the right-hand digit ratio and treatment expectancy added to Model (A1).*

*2F was built on Model 2B, with the addition of psychopathy scores and age.*

*2G includes the sex of the research assistant, sexual preference, number of children, and relationship status.*

*2H tests the influence of cortisol on cognitive empathy and its interaction with treatment using an interaction term specifying the binary treatment variable and continuous logged cortisol levels.*

Results showed no effect of testosterone treatment on the post-administration RMET score in any model, and no evidence for a moderating role of the digit ratios. Model (A1) beta = 0.267, 95% CI = [−0.40, 1.02], $t(237) = 0.37$, $P = 0.49$. The estimated effect size (Cohen's *d*) was 0.07, CI = [-0.13, 0.27].

To be consistent across experiments, we applied the following exclusion criteria: recent evidence indicates that similar to other commercially available ELISA-based kit, DRG's ELISA kit overestimates salivary testosterone levels by approximately 50% compared to LCMS (Welker et al. 2016). Therefore, for Experiment 2, our cutoff for 'high' salivary testosterone concentrations was adjusted to 600 pg/ml (400 pg/ml [from Experiment 1] x 1.50) to reflect the relatively higher salivary testosterone concentrations estimated using the DRG ELISA kit. We found 56 instances of testosterone levels in excess of the cutoff criterion at baseline (*high_pre_T* variable), 25 from the testosterone group and 31 in the Placebo group. Results are robust to exclusion of these participants (all *P*-values for treatment binary variable > 0.088).

Finally, the RMET score of a participant was greater than 4.5 SDs below the mean (cutoff threshold = 8.01; participant score = 8); all of the results are robust to his exclusion from the analysis (see Table S3b).

**Regressions Analyses: Testing for Effects of 2D:4D on Total RMET Performance**
**Experiment 1.** We estimate several regression models with the total RMET scores (morning + afternoon) as the DV and levels, and 2D:4D and controls as IVs for both studies:

*1J tests the effects of right hand 2D:4D on RMET performance.*

*1K tests the left 2D:4D.*

*1L tests the average 2D:4D.*

*1M tests for interaction effects between right hand 2D:4D and treatment binary*

*1N tests for interaction effects between left hand 2D:4D and treatment binary*

We find no significant relationships between RMET scores and 2D:4D or interactions with testosterone treatment (see Table S4a).

**Experiment 2**. We estimate several regression models with the total RMET scores (morning + afternoon) as the DV and levels, and 2D:4D and controls as IVs for both studies:

*2J tests the effects of right hand 2D:4D on RMET performance.*

*2K tests the left 2D:4D.*

*2L tests the average 2D:4D.*

*2M tests for interaction effects between right hand 2D:4D and treatment binary*

*2N tests for interaction effects between left hand 2D:4D and treatment binary*

We find no significant relationships between RMET scores and 2D:4D or interactions with testosterone treatment (Table S4b).

**Behavioral Results at the Question Level.** We further tested whether exogenous testosterone affected responses to specific RMET questions, by conducting Chi-squared tests that compared post-treatment responses between the testosterone and placebo groups.

**Experiment 1.** Consistent with an absence of signal in the data, only 2 out of the 36 uncorrected $P$-values (0.055%) were significant at the 5% level and none survived correction for multiple comparisons (initial alpha level $P$ = 0.05, Bonferroni corrected threshold $P$ = 0.05/36 = 0.001) (see Table S5a).

**Experiment 2.** Similarly, with an absence of signal in the data, only one question in Experiment 2 was significant yet did not survive correction for multiple comparisons (Initial alpha level $P$ = 0.05, Bonferroni corrected threshold $P$ = 0.05/36 = 0.001) (see Table S5b).

*References*

Carré, Justin M., Triana L. Ortiz, Brandy Labine, Benjamin J. P. Moreau, Essi Viding, Craig S. Neumann, and Bernard Goldfarb. 2015. "Digit Ratio (2D:4D) and Psychopathic Traits Moderate the Effect of Exogenous Testosterone on Socio-Cognitive Processes in Men." *Psychoneuroendocrinology* 62 (December): 319–26.

Eisenegger, Christoph, Arnold von Eckardstein, Ernst Fehr, and Sigrid von Eckardstein. 2013. "Pharmacokinetics of Testosterone and Estradiol Gel Preparations in Healthy Young Men." *Psychoneuroendocrinology* 38 (2): 171–78.

Frederick, Shane. 2005. "Cognitive Reflection and Decision Making." *The Journal of Economic Perspectives: A Journal of the American Economic Association* 19 (4): 25–42.

Gneezy, Uri, and Jan Potters. 1997. "An Experiment on Risk Taking and Evaluation Periods." *The Quarterly Journal of Economics* 112 (2): 631–45.

Henderson, Austin, Garrett Thoelen, Amos Nadler, Jorge Barraza, and Gideon Nave. 2018. "Testing the Influence of Testosterone Administration on Men's Honesty in a Large Laboratory Experiment." *Scientific Reports* 8 (1): 11556.

Holt, Charles A., and Susan K. Laury. 2005. "Risk Aversion and Incentive Effects: New Data without Order Effects." *The American Economic Review* 95 (3): 902–12.

Honk, Jack van, Dennis J. Schutter, Peter A. Bos, Anne-Wil Kruijt, Eef G. Lentjes, and Simon Baron-Cohen. 2011. "Testosterone Administration Impairs Cognitive Empathy in Women Depending on Second-to-Fourth Digit Ratio." *Proceedings of the National Academy of Sciences of the United States of America* 108 (8): 3448–52.

Horton, R., and J. F. Tait. 1966. "Androstenedione Production and Interconversion Rates Measured in Peripheral Blood and Studies on the Possible Site of Its Conversion to Testosterone." *The Journal of Clinical Investigation* 45 (3): 301–13.

Hucklebridge, F., T. Hussain, P. Evans, and A. Clow. 2005. "The Diurnal Patterns of the Adrenal Steroids Cortisol and Dehydroepiandrosterone (DHEA) in Relation to Awakening." *Psychoneuroendocrinology* 30 (1): 51–57.

Hurwitz, Shelley, Richard J. Cohen, and Gordon H. Williams. 2004. "Diurnal Variation of Aldosterone and Plasma Renin Activity: Timing Relation to Melatonin and Cortisol and Consistency after Prolonged Bed Rest." *Journal of Applied Physiology* 96 (4): 1406–14.

Kocoska-Maras, Ljiljana, Niklas Zethraeus, Angelique Flöter Rådestad, Tore Ellingsen, Bo von Schoultz, Magnus Johannesson, and Angelica Lindén Hirschberg. 2011. "A Randomized Trial of the Effect of Testosterone and Estrogen on Verbal Fluency, Verbal Memory, and Spatial Ability in Healthy Postmenopausal Women." *Fertility and Sterility* 95 (1): 152–57.

Manning, John T. 2011. "Resolving the Role of Prenatal Sex Steroids in the Development of Digit Ratio." *Proceedings of the National Academy of Sciences of the United States of America* 108 (39): 16143–44.

Nave, G., A. Nadler, D. Dubois, D. Zava, C. Camerer, and H. Plassmann. 2018. "Single-Dose Testosterone Administration Increases Men's Preference for Status Goods." *Nature Communications* 9 (1): 2433.

Niederle, Muriel, and Lise Vesterlund. 2005. "Do Women Shy Away From Competition? Do Men Compete Too Much?" Cambridge, MA: National Bureau of Economic Research. https://doi.org/10.3386/w11474.

Rogol, A. D., N. Tkachenko, and N. Bryson. 2016. "Natesto™ , a Novel Testosterone Nasal Gel, Normalizes Androgen Levels in Hypogonadal Men." *Andrology* 4 (1): 46–54.
Rolf, C., U. Knie, G. Lemmnitz, and E. Nieschlag. 2002. "Interpersonal Testosterone Transfer after Topical Application of a Newly Developed Testosterone Gel Preparation." *Clinical Endocrinology* 56 (5): 637–41.
Salameh, Wael A., Mildred M. Redor-Goldman, Nigel J. Clarke, Richard E. Reitz, and Michael P. Caulfield. 2010. "Validation of a Total Testosterone Assay Using High-Turbulence Liquid Chromatography Tandem Mass Spectrometry: Total and Free Testosterone Reference Ranges." *Steroids* 75 (2): 169–75.
Xu, Li, and David C. Spink. 2008. "Analysis of Steroidal Estrogens as Pyridine-3-Sulfonyl Derivatives by Liquid Chromatography Electrospray Tandem Mass Spectrometry." *Analytical Biochemistry* 375 (1): 105–14.
Zethraeus, Niklas, Ljiljana Kocoska-Maras, Tore Ellingsen, Bo von Schoultz, Angelica Lindén Hirschberg, and Magnus Johannesson. 2009. "A Randomized Trial of the Effect of Estrogen and Testosterone on Economic Behavior." *Proceedings of the National Academy of Sciences of the United States of America* 106 (16): 6535–38.

**Figure S1a. Experiment 1: Flow diagram of the progress through the phases of a parallel randomized trial of testosterone and placebo groups**

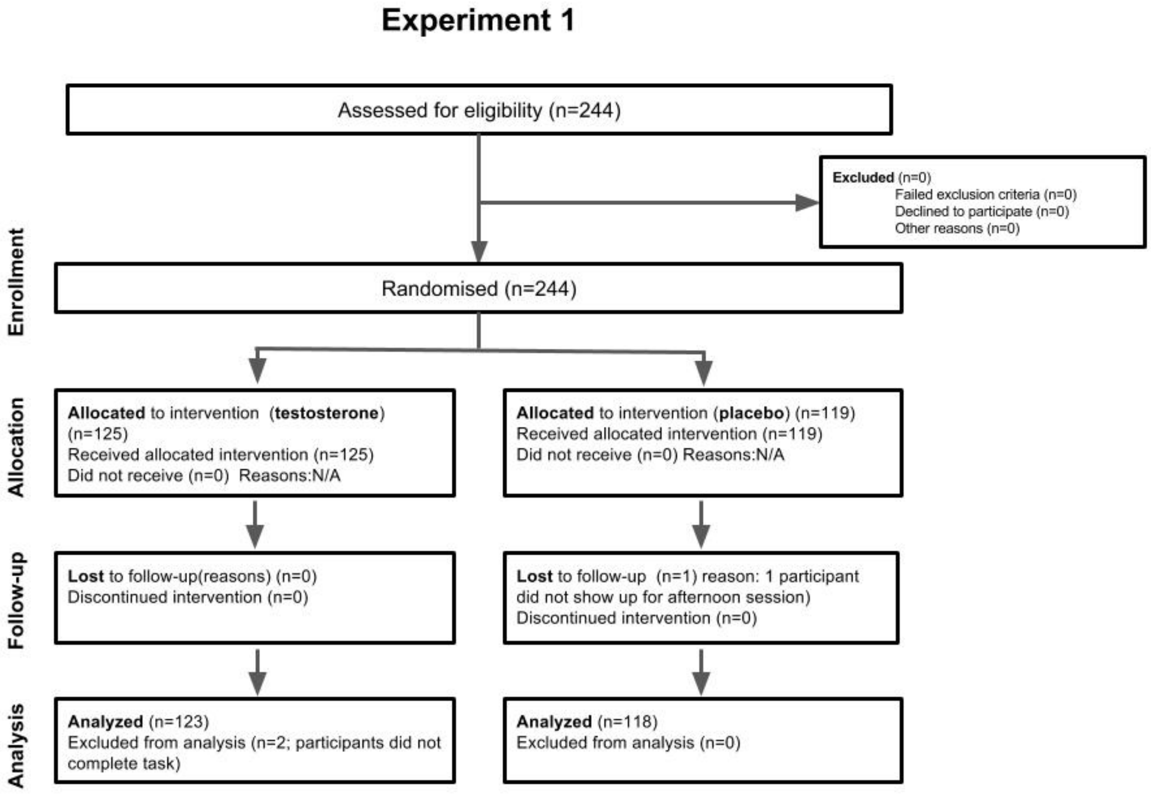

**The pathway of recruiting for Experiment 1 from eligibility assessment, randomization, allocation, and study execution are depicted with associated samples size**

**Figure S1b. Experiment 2: Flow diagram of the progress through the phases of a parallel randomized trial of testosterone and placebo groups**

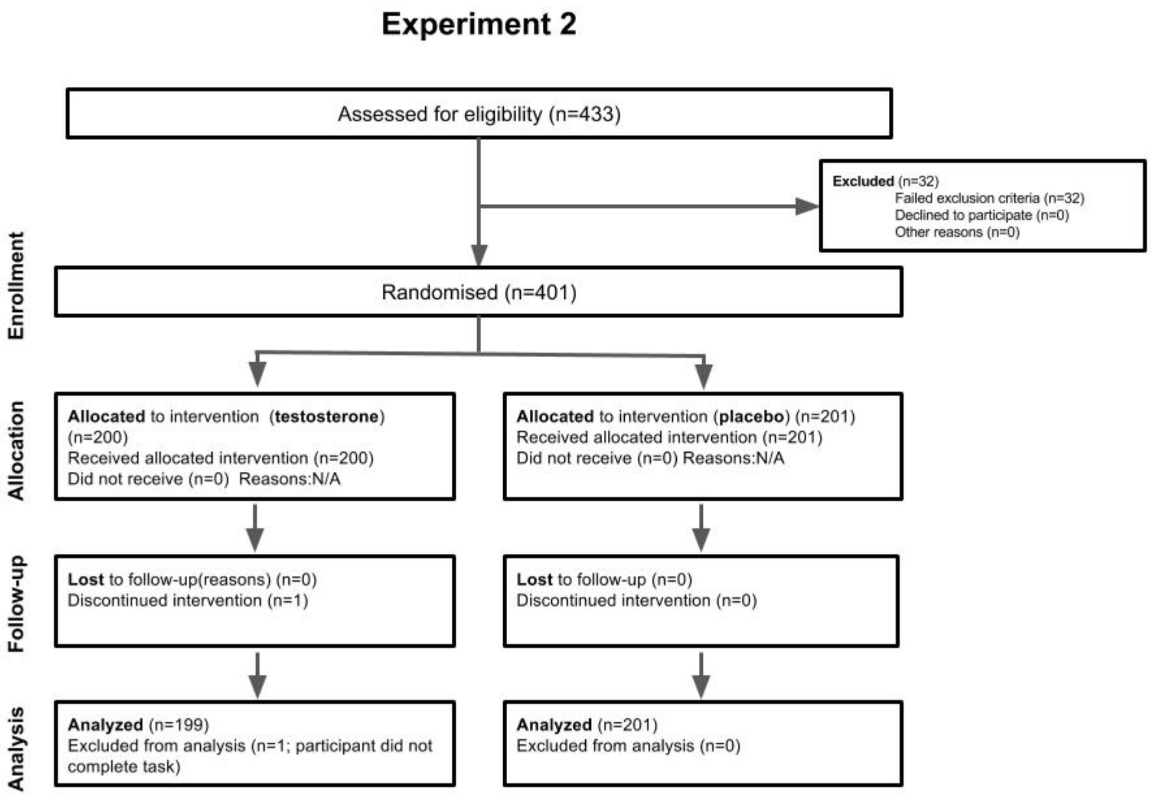

**The pathway of recruiting for Experiment 2 from eligibility assessment, randomization, allocation, and study execution are depicted with associated samples sizes.**

**Figure S2: Example RMET item**

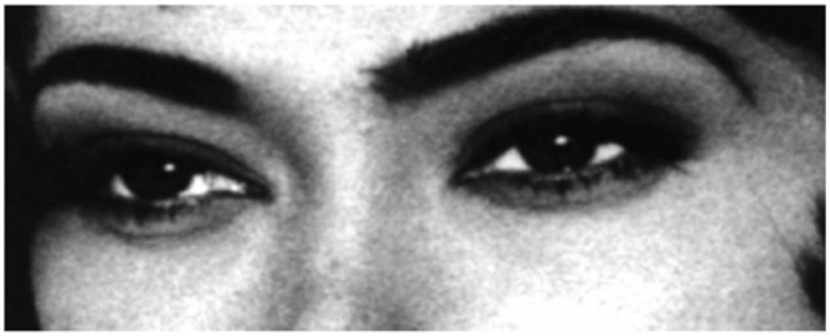

- aghast
- fantasizing
- impatient
- alarmed

**AGHAST**
**horrified, astonished, alarmed**
**Jane was aghast when she discovered her house had been burgled.**

**FANTASIZING**
**daydreaming**
**Emma was fantasizing about being a film star.**

**IMPATIENT**
**restless, wanting something to happen soon**
**Jane grew increasingly impatient as she waited for her friend who was already 20 minutes late.**

**ALARMED**
**fearful, worried, filled with anxiety**
**Claire was alarmed when she thought she was being followed home.**

**Fig. S2: Example item from the RMET as presented in the task, which includes a dictionary for each option to accommodate a potentially wide range of English proficiency among participants.**

**Figure S3: Pharmacokinetics of Single Dose Natesto® for Hypogonadal Men**

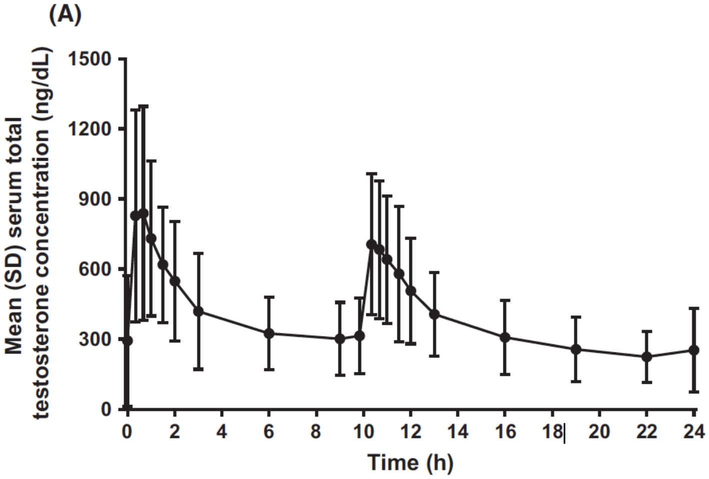

**Fig. S3: This figure depicts the pharmacokinetics of a single dose of Natesto® in men with below-normal testosterone levels in Rogol, Tkachenko, and Bryson (2016).**

.

**Figure S4. Effect of Testosterone Administration on RMET Scores in Experiment 1**

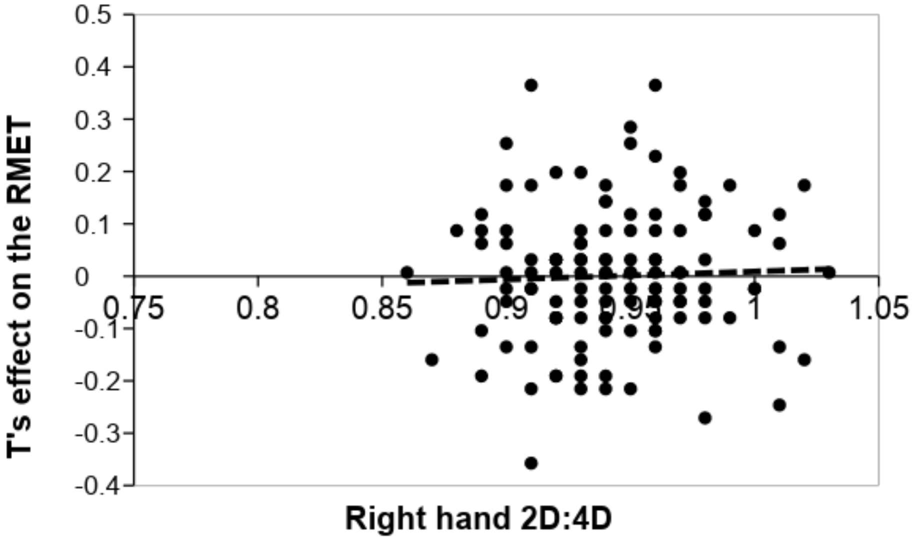

Fig. S4. No association was found between the 2D:4D and the effect of testosterone treatments on RMET scores (calculated as the normalized difference between the post-treatment and baseline RMET score in the testosterone group, after regressing out the effect of portion (A or B) order. The finding is robust to inclusion/exclusion of the latter control.

**Table S1a. Experiment 1: Self-reported demographic data summary (standard errors in parentheses)**

| | All | Testosterone | Placebo | *P*-values for t-test of difference |
|---|---|---|---|---|
| *n* | 243 | 118 | 125 | |
| Age | 23.63<br>(0.46) | 24.42<br>(0.77) | 22.78<br>(0.49) | 0.08 |
| Left-handed (proportion) | 0.074<br>(0.02) | 0.064<br>(0.02) | 0.085<br>(0.03) | 0.54 |
| Heterosexual (proportion) | 0.9<br>(0.02) | 0.91<br>(0.03) | 0.89<br>(0.03) | 0.56 |
| Treatment expectancy[1] | 2.76<br>(0.06) | 2.67<br>(0.08) | 2.85<br>(0.09) | 0.16 |
| Married (proportion) | 0.08<br>(0.02) | 0.09<br>(0.03) | 0.08<br>(0.03) | 0.74 |
| In a relationship (proportion) | 0.38<br>(0.03) | 0.34<br>(0.05) | 0.42<br>(0.04) | 0.2 |
| Has children | 0.06<br>(0.02) | 0.08<br>(0.02) | 0.04<br>(0.02) | 0.23 |
| Personal monthly income[2] | 2.05<br>(0.11) | 2.02<br>(0.14) | 2.07<br>(0.16) | 0.84 |
| Positive affect[3] | 2.61<br>(0.06) | 2.63<br>(0.04) | 2.6<br>(0.09) | 0.78 |
| Negative affect[3] | 1.45<br>(0.04) | 1.46<br>(0.05) | 1.43<br>(0.05) | 0.93 |

1 5-point scale, 1 = definitely did not receive testosterone, 2 = probably did not, 3 = unsure, 4 = probably received testosterone, 5 = definitely received testosterone

2 5-point scale, 1 = < $500/month, 2 = $501–$1,000/month, 3 = $1,001–$1,500/month, 4 = $1,501–$2,000/month 5 = > $2001/month

3 These affective scores were taken in afternoon (post-treatment)

**Table S1b. Experiment 2: Self-reported demographic data summary (standard errors in parentheses)**

| | All | Testosterone | Placebo | *P*-values for t-test of difference |
|---|---|---|---|---|
| *n* | 400 | 199 | 201 | |
| Age | 22.8<br>(0.24) | 23.06<br>(0.35) | 22.55<br>(0.32) | 0.28 |
| Heterosexual (proportion) | 0.94<br>(0.01) | 0.94<br>(0.02) | 0.94<br>(0.02) | 0.91 |
| Treatment expectancy[1] | 1.7<br>(0.02) | 1.73<br>(0.03) | 1.67<br>(0.03) | 0.25 |
| Married (proportion) | 0.05<br>(0.01) | 0.07<br>(0.02) | 0.04<br>(0.01) | 0.11 |
| In a relationship (proportion) | 0.49<br>(0.03) | 0.47<br>(0.04) | 0.51<br>(0.04) | 0.45 |
| Has children | 0.11<br>(0.02) | 0.13<br>(0.02) | 0.09<br>(0.02) | 0.26 |
| Personal yearly income[2] | 1.31<br>(0.03) | 1.36<br>(0.05) | 1.27<br>(0.04) | 0.19 |
| Psychopathy Factor 1 | 33.53<br>(0.48) | 33.45<br>(0.67) | 33.73<br>(0.69) | 0.78 |
| Psychopathy Factor 2 | 31.95<br>(0.45) | 32.09<br>(0.63) | 31.96<br>(0.64) | 0.88 |

1 1 = thought that they received testosterone, 2 = thought that they received placebo.
2 6-point scale, 1 = < $20,000/year, 2 = $20,000-40,000/year, 3 = $40,000-60,000/year, 4 = $60,000-80,000/year, 5 =$80,000-100,000/year, 6 = > $100,000/year

**Table S2a. Experiment 1: Hormone panel measurements pg/mL summary statistics (standard errors in parentheses)**

| | Placebo | | Testosterone | | Two-tailed p-value from t-test of T-Placebo equality | |
|---|---|---|---|---|---|---|
| **Sampling time**[1] | **9:00 AM** | **2:00 PM** | **9:00 AM** | **2:00 PM** | **9:00 AM** | **2:00 PM** |
| Testosterone | 613 | 278 | 486 | 11,449 | 0.30 | < 0.0001 |
| | (96.6) | (37.5) | (75.0) | (1,386) | | |
| Androstenedione | 101.1 | 75.0 | 99.9 | 380.1 | 0.83 | < 0.0001 |
| | (4.82) | (3.35) | (3.32) | (48.9) | | |
| DHT | 11.59 | 8.61 | 8.56 | 79.2 | 0.19 | < 0.0001 |
| | (2.11) | (1.00) | (1.00) | (14.48) | | |
| Progesterone | 9.65 | 10.9 | 8.21 | 9.18 | 0.38 | 0.43 |
| | (1.40) | (1.73) | (0.90) | (1.34) | | |
| Progesterone170H | 29.6 | 16.9 | 29.7 | 14.5 | 0.97 | 0.05 |
| | (1.75) | (0.85) | (1.97) | (0.78) | | |
| Estrone | 1.13 | 0.75 | 1.28 | 0.81 | 0.23 | 0.42 |
| | (0.08) | (0.06) | (0.09) | (0.04) | | |
| Estradiol | 0.55 | 0.52 | 0.55 | 0.51 | 0.95 | 0.73 |
| | (0.03) | (0.03) | (0.03) | (0.02) | | |
| DHEA | 207 | 117 | 194 | 113 | 0.36 | 0.67 |
| | (10.3) | (6.44) | (9.34) | (5.80) | | |
| Deoxycortisol11 | 20.7 | 11.5 | 21.9 | 10.0 | 0.57 | 0.20 |
| | (1.28) | (0.88) | (1.66) | (0.74) | | |
| Cortisol | 3.61 | 1.35 | 3.44 | 1.32 | 0.68 | 0.81 |
| | (0.36) | (0.09) | (0.20) | (0.08) | | |
| Cortisone | 13.3 | 8.05 | 13.5 | 8.17 | 0.70 | 0.76 |
| | (0.37) | (0.30) | (0.44) | (0.25) | | |
| Corticosterone | 42.4 | 14.4 | 47.8 | 11.3 | 0.53 | 0.3 |
| | (5.53) | (2.64) | (6.57) | (1.62) | | |
| Aldosterone | 22.3 | 23.0 | 23.7 | 21.1 | 0.45 | 0.26 |
| | -1.19 | (1.20) | (1.30) | (1.04) | | |
| Melatonin | 9.83 | 8.48 | 25.4 | 18.1 | 0.39 | 0.53 |
| | (0.93) | (4.52) | (15.3) | (9.58) | | |

1: Main effects of time (afternoon vs. treatment) are due to the diurnal cycles of the hormones (Rowe et al. 1974; Brambilla et al. 2009)

**Table S2b. Experiment 2: Hormone panel measurements testosterone and cortisol pg/mL summary statistics (standard errors in parentheses)**

| | **Placebo** (N = 201) | | **Testosterone** (N = 199) | | Two-tailed p-value from t-test of T-Placebo equality | |
|---|---|---|---|---|---|---|
| | **Baseline** | **Post-treatment** | **Baseline** | **Post-treatment** | **Baseline** | **Post-treatment** |
| Testosterone | 324<br>(25.0) | 395<br>(25.3) | 318<br>(26.9) | 8,966<br>(1,044) | 0.87 | < 0.0001 |
| Cortisol | 3.21<br>(0.189) | | 3.28<br>(0.163) | | 0.80 | *na* |

**Table S3a. Experiment 1: OLS regressions, dependent variable: RMET afternoon**

| VARIABLES | 1A | 1B | 1C | 1D | 1E | 1F | 1G | 1H | 1I |
|---|---|---|---|---|---|---|---|---|---|
| Treatment | 0.106 | 8.641 | 6.052 | 9.001 | 0.154 | 8.798 | 8.976 | 9.197 | |
| | (0.286) | (8.070) | (8.306) | (9.029) | (0.313) | (8.148) | (8.068) | (8.264) | |
| Morning | 0.457*** | 0.461*** | 0.463*** | 0.465*** | 0.439*** | 0.441*** | 0.420*** | 0.426*** | 0.421*** |
| | (0.066) | (0.067) | (0.067) | (0.067) | (0.070) | (0.068) | (0.068) | (0.070) | (0.067) |
| Order | -0.478* | -0.507* | -0.479* | -0.487* | -0.491 | -0.639** | -0.596** | -0.590* | -0.582* |
| | (0.289) | (0.291) | (0.290) | (0.290) | (0.303) | (0.296) | (0.296) | (0.300) | (0.300) |
| Digit ratio (right) | | 8.519 | | | 3.921 | 8.695 | 9.248 | 9.345 | 4.332 |
| | | (6.017) | | | (4.420) | (5.983) | (5.928) | (6.052) | (4.279) |
| Digit ratio (left) | | | 1.468 | | | | | | |
| | | | (6.202) | | | | | | |
| Digit ratio (average) | | | | 6.297 | | | | | |
| | | | | (6.790) | | | | | |
| Treatment Expectancy | | | | | -0.0437 | 0.0911 | 0.0859 | 0.0783 | 0.0686 |
| | | | | | (0.159) | (0.160) | (0.159) | (0.162) | (0.161) |
| Estradiol | | | | | -0.0546 | | | | |
| | | | | | (0.229) | | | | |
| DHEA | | | | | 0.0478 | | | | |
| | | | | | (0.335) | | | | |
| Progesterone | | | | | 0.132 | | | | |
| | | | | | (0.245) | | | | |
| Progesterone 170H | | | | | -0.0100 | | | | |
| | | | | | (0.273) | | | | |
| Deoxycortisol11 | | | | | -0.0503 | | | | |
| | | | | | (0.279) | | | | |
| Cortisol | | | | | 0.561 | | | | |
| | | | | | (0.469) | | | | |
| Cortisone | | | | | -0.0297 | | | | |
| | | | | | (0.436) | | | | |
| Corticosterone | | | | | -0.172 | | | | |
| | | | | | (0.259) | | | | |
| Aldosterone | | | | | -0.0683 | | | | |
| | | | | | (0.205) | | | | |
| Melatonin | | | | | 0.0610 | | | | |
| | | | | | (0.321) | | | | |
| DHEA7 | | | | | 0.0626 | | | | |
| | | | | | (0.348) | | | | |
| Digit ratio (R) x Treatment | | -8.956 | | | | -9.100 | -9.177 | -9.413 | 0.293 |
| | | (8.506) | | | | (8.588) | (8.503) | (8.710) | (0.320) |
| Digit ratio (L) x Treatment | | | -6.291 | | | | | | |
| | | | (8.776) | | | | | | |
| Digit Ratio (A) x Treatment | | | | -9.383 | | | | | |
| | | | | (9.530) | | | | | |
| Negative Affect | | | | | | -0.366 | -0.363 | -0.376 | -0.386 |
| | | | | | | (0.282) | (0.279) | (0.288) | (0.287) |
| Positive Affect | | | | | | 0.0326 | 0.0381 | 0.0404 | 0.0108 |
| | | | | | | (0.157) | (0.158) | (0.161) | (0.160) |
| Age | | | | | | -0.0422** | -0.0367* | -0.0297 | -0.0312 |
| | | | | | | (0.0207) | (0.0210) | (0.0289) | (0.0287) |
| CRT | | | | | | | 0.258* | 0.269* | 0.278* |
| | | | | | | | (0.140) | (0.144) | (0.144) |
| Math | | | | | | | 0.0428 | 0.0442 | 0.0436 |

| | (1) | (2) | (3) | (4) | (5) | (6) | (7) | (8) | (9) |
|---|---|---|---|---|---|---|---|---|---|
| | | | | | | | (0.0382) | (0.0390) | (0.0389) |
| Heterosexual | | | | | | | | 0.0609 | 0.0490 |
| | | | | | | | | (0.279) | (0.279) |
| Married | | | | | | | | 0.188 | 0.152 |
| | | | | | | | | (0.772) | (0.772) |
| In a relationship | | | | | | | | 0.0882 | 0.126 |
| | | | | | | | | (0.342) | (0.342) |
| Sexual Experience | | | | | | | | -0.0767 | -0.0938 |
| | | | | | | | | (0.157) | (0.157) |
| Has children | | | | | | | | -0.00277 | 0.128 |
| | | | | | | | | (0.386) | (0.383) |
| 0b.treatment#cortisol | | | | | | | | | 0.518* |
| | | | | | | | | | (0.303) |
| 1.treatment#cortisol | | | | | | | | | 0.177 |
| | | | | | | | | | (0.315) |
| Constant | 7.912*** | -0.250 | 6.443 | 1.825 | 4.249 | 1.118 | -0.295 | -0.977 | 3.909 |
| | (0.905) | (5.841) | (6.052) | (6.594) | (4.737) | (5.903) | (5.871) | (6.258) | (4.674) |
| Observations | 241 | 240 | 241 | 241 | 238 | 228 | 228 | 228 | 228 |
| R-squared | 0.171 | 0.178 | 0.174 | 0.175 | 0.190 | 0.192 | 0.215 | 0.217 | 0.224 |

Standard errors in parentheses
*** $p < 0.01$, ** $p < 0.05$, * $p < 0.10$

### Table S3b. Experiment 2: OLS regressions, dependent variable: RMET Score

**Experiment 2 Data OLS Regressions with Total Score as DV**

| VARIABLES | 2A | 2B | 2C | 2D | 2E | 2F | 2G | 2G |
|---|---|---|---|---|---|---|---|---|
| Treatment | 0.244 | 20.64* | 16.54 | 23.00* | 0.177 | 18.04 | 21.58* | |
| | (0.385) | (11.94) | (11.66) | (13.18) | (0.381) | (11.54) | (11.87) | |
| Digit ratio (right) | | 12.78 | | | | 6.129 | 8.282 | -3.801 |
| | | (8.978) | | | | (8.699) | (8.821) | (6.329) |
| Digit ratio (left) | | | 10.29 | | | | | |
| | | | (8.736) | | | | | |
| Digit ratio (average) | | | | 14.00 | | | | |
| | | | | (9.770) | | | | |
| Digit ratio (R) x Treatment | | -21.47* | | | | -18.80 | -22.37* | 0.752 |
| | | (12.53) | | | | (12.11) | (12.45) | (0.730) |
| Digit ratio (L) x Treatment | | | -17.27 | | | | | |
| | | | (12.30) | | | | | |
| Digit Ratio (A) x Treatment | | | | -24.03* | | | | |
| | | | | (13.87) | | | | |
| Treatment Expectancy | | | | | 1.302*** | 1.185*** | 1.022** | 1.045** |
| | | | | | (0.415) | (0.435) | (0.448) | (0.454) |
| Factor 1 psychopathy | | | | | | -0.00812 | -0.0151 | -0.0159 |
| | | | | | | (0.0276) | (0.0287) | (0.0289) |
| Factor 2 psychopathy | | | | | | 0.0285 | 0.0296 | 0.0248 |
| | | | | | | (0.0300) | (0.0314) | (0.0316) |
| CRT | | | | | | 0.764*** | 0.666*** | 0.643*** |
| | | | | | | (0.180) | (0.186) | (0.188) |
| Sex of RA | | | | | | | 0.957** | 0.901* |
| | | | | | | | (0.481) | (0.484) |
| Age | | | | | | 0.120*** | 0.0793 | 0.0973* |
| | | | | | | (0.0442) | (0.0545) | (0.0560) |
| Heterosexual | | | | | | | -0.789 | -0.677 |
| | | | | | | | (0.829) | (0.835) |
| Married | | | | | | | 0.353 | 0.131 |
| | | | | | | | (1.045) | (1.058) |
| In a relationship | | | | | | | 0.723* | 0.648 |
| | | | | | | | (0.425) | (0.430) |
| Has children | | | | | | | -0.0533 | -0.0640 |
| | | | | | | | (0.391) | (0.395) |
| 0b.treatment#cortisol | | | | | | | | 0.0226 |
| | | | | | | | | (0.112) |
| 1.treatment#cortisol | | | | | | | | -0.131 |
| | | | | | | | | (0.128) |
| Constant | 25.64*** | 13.51 | 15.94* | 12.38 | 23.46*** | 13.73 | 12.27 | 23.54*** |
| | (0.270) | (8.573) | (8.291) | (9.299) | (0.745) | (8.426) | (8.649) | (6.315) |
| Observations | 398 | 355 | 354 | 354 | 398 | 349 | 332 | 329 |
| R-squared | 0.001 | 0.009 | 0.006 | 0.009 | 0.025 | 0.100 | 0.109 | 0.105 |

Standard errors in parentheses

*** p<0.01, ** p<0.05, * p<0.1

**Table S4a. Experiment 1: OLS regressions with total RMET score as DV**

**Influence of Digit Ratio and Treatment on Total Score**

| VARIABLES | 1J | 1K | 1L | 1M | 1N |
|---|---|---|---|---|---|
| Digit ratio (right) | 3.659 | | | | |
| | -7.312 | | | | |
| Digit ratio (left) | | -4.737 | | | |
| | | -7.545 | | | |
| Digit ratio (average) | | | -0.386 | | |
| | | | -8.191 | | |
| Placebo Treatment x Digit Ratio (R) Interaction | | | | 4.488 | |
| | | | | -7.352 | |
| T Treatment x Digit Ratio (R) Interaction | | | | 5.05 | |
| | | | | -7.427 | |
| Placebo Treatment x Digit Ratio (L) Interaction | | | | | -4.695 |
| | | | | | -7.547 |
| T Treatment x Digit Ratio (L) Interaction | | | | | -4.198 |
| | | | | | -7.568 |
| Constant | 24.06*** | 32.00*** | 27.89*** | 23.01*** | 31.72*** |
| | -6.937 | -7.142 | -7.761 | -7.006 | -7.15 |
| Observations | 240 | 241 | 241 | 240 | 241 |
| R-squared | 0.001 | 0.002 | 0 | 0.006 | 0.005 |

Standard errors in parentheses
*** *p* < 0.01, ** *p* < 0.05, * *p* < 0.10

### Table S4b. Experiment 2: OLS regressions with total RMET score as DV

**Influence of Digit Ratio and Treatment on Total Score**

| VARIABLES | 2J | 2K | 2L | 2M | 2N |
|---|---|---|---|---|---|
| Digit ratio (right) | -0.779 | | | | |
| | -6.411 | | | | |
| Digit ratio (left) | | 1.309 | | | |
| | | -6.316 | | | |
| Digit ratio (average) | | | 0.363 | | |
| | | | -7.119 | | |
| Placebo Treatment x Digit Ratio (R) Interaction | | | | -0.767 | |
| | | | | -6.42 | |
| Testosterone Treatment x Digit Ratio (R) Interaction | | | | -0.69 | |
| | | | | -6.44 | |
| Placebo Treatment x Digit Ratio (L) Interaction | | | | | 1.317 |
| | | | | | -6.325 |
| Testosterone Treatment x Digit Ratio (L) Interaction | | | | | 1.407 |
| | | | | | -6.343 |
| Constant | 26.48*** | 24.50*** | 25.39*** | 26.43*** | 24.45*** |
| | -6.11 | -5.984 | -6.764 | -6.124 | -5.997 |
| Observations | 356 | 355 | 355 | 356 | 355 |
| R-squared | 0 | 0 | 0 | 0 | 0 |

Standard errors in parentheses
*** $p < 0.01$, ** $p < 0.05$, * $p < 0.10$

**Table S5a: Experiment 1: Post-treatment comparison of scores by Chi-squared tests for each question**

| Group B (completed questions 19–36 first), *N* = 123 | | | Group A (completed questions 1–18 first), *N* = 118 | | |
|---|---|---|---|---|---|
| Item | Chi-squared | *P*-value (uncorrected) | Item | Chi-squared | *P*-value (uncorrected) |
| 1 | 7.99 | 0.05 | 19 | 4.33 | 0.23 |
| 2 | 0.30 | 0.94 | 20 | 3.95 | 0.27 |
| 3 | 7.33 | 0.06 | 21 | 1.22 | 0.54 |
| 4 | 0.33 | 0.85 | 22 | 4.97 | 0.08 |
| 5 | 6.14 | 0.11 | 23 | 5.77 | 0.12 |
| 6 | 5.19 | 0.15 | 24 | 3.00 | 0.39 |
| 7 | 8.90 | 0.03 | 25 | 4.92 | 0.18 |
| 8 | 1.67 | 0.64 | 26 | 1.56 | 0.66 |
| 9 | 3.40 | 0.33 | 27 | 4.36 | 0.11 |
| 10 | 2.33 | 0.51 | 28 | 2.81 | 0.42 |
| 11 | 4.96 | 0.17 | 29 | 3.25 | 0.35 |
| 12 | 3.87 | 0.28 | 30 | 2.51 | 0.47 |
| 13 | 4.34 | 0.23 | 31 | 2.55 | 0.47 |
| 14 | 0.66 | 0.72 | 32 | 1.25 | 0.74 |
| 15 | 1.94 | 0.59 | 33 | 3.82 | 0.28 |
| 16 | 5.95 | 0.11 | 34 | 4.99 | 0.17 |
| 17 | 0.86 | 0.84 | 35 | 2.89 | 0.41 |
| 18 | 1.76 | 0.62 | 36 | 2.61 | 0.46 |

**Table S5b: Experiment 2: Post-treatment comparison of scores by Chi-squared tests for each question**

| Item | Chi-squared | *P*-value (uncorrected) | Item | Chi-squared | *P*-value (uncorrected) |
|---|---|---|---|---|---|
| 1 | 0.07 | 0.786 | 19 | 0.02 | 0.9 |
| 2 | 1.09 | 0.296 | 20 | 0.26 | 0.61 |
| 3 | 0.09 | 0.766 | 21 | 0.24 | 0.62 |
| 4 | 0.28 | 0.598 | 22 | 1.46 | 0.23 |
| 5 | 0.35 | 0.553 | 23 | 3.42 | 0.06 |
| 6 | 3.43 | 0.064 | 24 | 0.65 | 0.42 |
| 7 | 4.62 | 0.032 | 25 | 0.05 | 0.82 |
| 8 | 0.01 | 0.935 | 26 | 2.03 | 0.15 |
| 9 | 1.53 | 0.216 | 27 | 1.35 | 0.25 |
| 10 | 0.18 | 0.669 | 28 | 0.52 | 0.47 |
| 11 | 1.51 | 0.22 | 29 | 3.05 | 0.08 |
| 12 | 0.00 | 0.965 | 30 | 0.2 | 0.65 |
| 13 | 3.07 | 0.08 | 31 | 1.68 | 0.2 |
| 14 | 2.67 | 1.02 | 32 | 5.05 | 0.03 |
| 15 | 2.09 | 0.148 | 33 | 0 | 1 |
| 16 | 0.09 | 0.765 | 34 | 0.05 | 0.83 |
| 17 | 0.28 | 0.596 | 35 | 0.18 | 0.67 |
| 18 | 1.48 | 0.224 | 36 | 0.23 | 0.63 |

**Table S6a. Experiment 1: Correlations among all 2D:4D measures (*P*-values shown below)**

| | Right-hand 2D:4D | Left-hand 2D:4D | Average of right- and left-hand 2D:4D |
|---|---|---|---|
| Right-hand 2D:4D | 1 | | |
| Left-hand 2D:4D | 0.638 | 1 | |
| | 0.000 | | |
| Average of right- and left hand 2D:4D | 0.899 | 0.899 | 1 |
| | 0.000 | 0.000 | |

**Table S6b. Experiment 2: Correlations among all 2D:4D measures (*P*-values shown below)**

| | Right-hand 2D:4D | Left-hand 2D:4D | Average of right- and left-hand 2D:4D |
|---|---|---|---|
| Right-hand 2D:4D | 1 | | |
| Left-hand 2D:4D | 0.599 | 1 | |
| | 0.00 | | |
| Average of right- and left hand 2D:4D | 0.898 | 0.896 | 1 |
| | 0.00 | 0.00 | |

**Table S7a. Experiment 1: Detection levels, precision and normality tests of hormonal assay**

| Analyte | LLOQ | Range | Precision | Proportion undetected, pre-treatment sample A | Proportion undetected, first post-treatment sample B | K-S test *P*-value | K-S test (log) *P*-value |
|---|---|---|---|---|---|---|---|
| Estrone pg/mL | 0.5 | 0.5–510 | 8.7–13.7% | 0.132 | 0.257 | <0.01 | 0.56 |
| Estradiol pg/mL | 0.3 | 0.3–510 | 4.3–18.7% | 0.12 | 0.329 | 0.06 | 0.88 |
| Testosterone pg/mL | 3.0 | 3.0–5100 | 3.0–18.1% | 0 | 0.008 | $<10^{-20}$ | < 0.01 |
| Androstenedion e pg/mL | 5.0 | 5.0–2300 | 5.2–6.6% | 0 | 0.008 | $<10^{-20}$ | 0.008 |
| DHEA pg/mL | 20.0 | 20.0–1800 | 4.1–15.2% | 0.04 | 0.012 | 0.002 | 0.98 |
| DHT pg/mL | 10.0 | 10.0–920 | 3.6–17.7% | 0.786 | 0.473 | $<10^{-11}$ | 0.02 |
| Progesterone pg/mL | 10.0 | 10.0–10000 | 4.8–10.8% | 0.794 | 0.753 | <0.01 | 0.03 |
| 17OH-Progesterone pg/mL | 5.0 | 5.0–630 | 3.9–13.8% | 0.004 | 0.061 | 0.003 | 0.98 |
| 11-Deoxycortisol pg/mL | 5.0 | 5.0–410 | 6.8–16.6% | 0.132 | 0.473 | <0.01 | 0.04 |
| Cortisol ng/mL | 0.1 | 0.1–52 | 5.1–17.9% | 0 | 0.008 | <0.01 | 0.92 |
| Cortisone ng/mL | 0.1 | 0.1–81 | 4.1–14.9% | 0 | 0.008 | 0.07 | 0.59 |
| Corticosterone pg/mL | 5.0 | 5.0–1800 | 4.6–17.5% | 0.313 | 0.312 | <0.01 | 0.08 |
| Aldosterone pg/mL | 10.0 | 10.0–560 | 8.9–18.8% | 0.272 | 0.272 | <0.06 | 0.39 |
| Melatonin pg/mL | 2.5 | 2.5–10000 | 5.2–15.9% | 0.502 | 0.500 | 0.07 | 0.14 |

Note: *P*-values are calculated using a Kolmogorov-Smirnov test for the distributions of the second saliva sample compared with a Gaussian distribution, and for the log-transform (the null hypothesis is normal Gaussian distribution).

**Table S7b. Experiment 2: Detection levels, precision and normality tests of hormonal assay**

| Analyte | LLOQ | Range | Precision | Proportion undetected, pre-treatment sample A | Proportion undetected, first post-treatment sample B | K-S test *P*-value | K-S test (log) *P*-value | S-W test *P*-value | S-W test (log) *P*-value |
|---|---|---|---|---|---|---|---|---|---|
| Testosterone pg/mL | 10.07 | 0.94 - 1000 | 8% | 0 | 0 | $< 10^{-20}$ | < 0.01 | $< 10^{-5}$ | $< 10^{-5}$ |
| Cortisol ng/mL | 0.15 | 0.09 - 30 | 8.1% | 0 | 0 | < 0.01 | 0.92 | $< 10^{-5}$ | 0.29 |

**Additional datasets S1 and S2 (provided in separate files)**

Datasets containing participant ID numbers, RMET scores, survey responses, and hormonal measures are available in these files.
